# Predictability of ecological and evolutionary dynamics in a changing world

**DOI:** 10.1101/2023.11.01.565089

**Authors:** Claudio Bozzuto, Anthony R. Ives

## Abstract

Ecological and evolutionary predictions are being increasingly employed to inform decision-makers confronted with intensifying pressures menacing life on Earth. For these efforts to effectively guide conservation actions, knowing the limit of predictability is pivotal. In this study, we provide realistic expectations about the enterprise of predicting changes in ecological and evolutionary observations through time. We begin with an intuitive explanation of predictability (that is, the extent to which predictions are possible) employing an easy-to-use metric, predictive power *PP*(*t*). To illustrate the challenge of forecasting, we then show that among insects, birds, fishes, and mammals (i) 50% of the populations are predictable at most one year in advance, and (ii) the median one-year-ahead predictive power corresponds to a sobering prediction *R*^2^ of approximately 20%. Nonetheless, predictability is not an immutable property of ecological systems. For example, different harvesting strategies can impact the predictability of exploited populations to varying degrees. Moreover, considering multivariate time series, incorporating explanatory variables or accounting for time trends (environmental forcing) can enhance predictability. To effectively address the urgent challenge of biodiversity loss, researchers and practitioners must be aware of the predictive information within the available data and explore efficient ways to leverage this information for environmental stewardship.

## 1. Introduction

Three decades ago, Jared Diamond [1] and Edward O. Wilson [2] personified the main human-caused stressors to biodiversity as an evil quartet and the four mindless horsemen of the environmental apocalypse: habitat degradation, fragmentation and loss; overexploitation; introduction of non-native species and diseases; and pollution. Not only do these threats continue to impact life on Earth, they are joined by global warming causing and exacerbating the decline of biodiversity worldwide [3,4]. Worryingly, these stressors combine to form a perfect storm because they can act synergistically [5], and their impacts can percolate via cascading effects through entire ecosystems: biodiversity loss may be regarded as a harbinger of ecosystem collapse [6]. In addition to threatening population persistence and species richness, human stressors are potent agents of rapid evolutionary change in the wild, with exploitation leading the other stressors in driving phenotypic changes in populations [7,8]. The current geological epoch has been named the Anthropocene, one aspect of which is a human-caused “mutilation of the tree of life” [9], with “catastrophic and ongoing loss” of amphibian diversity [10] and insects dying “by a thousand cuts” [11].

Our understanding of the mechanisms underlying stressor effects and the global extent of these effects on biodiversity has improved (e.g. [12–14]), and collections of long-term data allow testing hypotheses about wildlife population changes (e.g. [15] and references therein). Global synoptic indicators of biodiversity change include the Red List Index [16] and the Living Planet Index [17]. These indicators are invaluable to inform – and ideally prompt actions by – governments and the general public [18]. Several authors have nonetheless bemoaned conservation biology and restoration ecology as generally lacking substantial efforts to model and predict future changes to guide mitigation actions (e.g. [19–21]). The reasons are multifaceted, and mechanistically predicting biodiversity responses to current and predicted future stressor levels continues to be a formidable task (e.g. [22]; but see [23]). Yet, Bodner *et al*. ([24], p.10) aptly note that “as researchers, we are often acutely aware of how much we do not know and therefore get stuck at ‘more research is required’. However, environmental changes are increasingly affecting our world, and decisions are made whether or not we are involved.”

Compared to the ecological dimensions of biodiversity loss, our understanding of the global distribution and the extent of human-induced evolutionary changes and genetic diversity loss is less developed [12,25]. This is unfortunate because genetic variation is the raw material enabling species to robustly adapt to a changing world [25]. Despite the growing recognition of this dimension of biodiversity, predictions in evolutionary science and applications face many challenges similar to ecologically oriented disciplines [26–28]. Evolutionary changes depend on changes in the genetic make-up of populations through time, but long-term genetic datasets continue to be rare. Therefore, analyses are often performed on phenotypic changes in the wild, which are routinely measured [13]. Phenotypic changes, however, do not necessarily imply genetic changes; instead, they could be ‘simply’ the result of phenotypic plasticity. It is generally challenging to discern plastic from genetic changes: Hendry *et al*. [13] expect genetic change to be more important for larger populations, for shorter generation times, for novel environmental conditions, and for pronounced environmental changes. Phenotypic changes are not sufficient to conclude there are genotypic changes, but phenotypic changes are necessarily observable for traits that are evolving. Therefore, in the absence of long-term genetic datasets, phenotypic datasets should serve in their stead.

How well can future population changes in size or phenotype be predicted? And is it fundamentally possible to predict future states and changes of biodiversity [29]? Answers to the first question often center around forecast accuracy (e.g. [30]), that is, how good a model has to be to make good predictions. Unfortunately, even a very good model can fail to make useful predictions of inherently unpredictable events. For example, assuming heads and tails have equal probability is an excellent statistical model for flipping a coin, but this is not much help in predicting the outcome of the next flip. Therefore, instead of focusing on how well models fit data, we address the second question by asking how much predictive information is contained in a dataset in the first place, and how a lack of information determines a barrier to prediction, dictated by a predictability limit of the system [29–33]. Here, we follow the definition that “predictability is the study of the extent to which predictions are possible” [32, p. 2425]. While the issue of an inherent limit of predictability has been studied for several decades in different fields, chief among them climatology [34], ecologists and evolutionary biologists have only recently started tackling it [26,27,29,30,35].

Our overall goal is to argue that biodiversity conservation needs to adopt the concept of predictability to better design strategies and policies to constrain human-caused threats to life on Earth. After an intuitive presentation of the concept of predictability, we apply it to approximately 1,600 ecological and phenotypic time series of invertebrate and vertebrate species to make plain how unpredictable ecological and evolutionary systems appear to be. We further discuss predictability addressing two central topics in biodiversity conservation: exploitation and time trends in abundance and phenotypic traits. Along the way, we make clear that predictability is not an immutable property of ecological systems but instead can be influenced, for example, by the way in which humans influence systems. Furthermore, we offer statistical evidence and theoretical proof for several previously stated but unresolved hypotheses ([30,36], and references therein). We conclude by summarizing why adopting the concept of predictability in biodiversity conservation will improve our strategies to counteract human-caused stressors of wildlife populations.

## 2. What is predictability?

### 2.1 An intuitive introduction

Instead of an exhaustive technical description of predictability (see e.g. [30–32]), we introduce the concept using an illustrative ecological example, the population of wolves on Isle Royale (Michigan, USA). We focus on the period 1959 – 2011 when the population experienced large fluctuations, but before it started a precipitous decline [37] (Fig. 1, ESM Appendix 2).

**Fig. 1.**
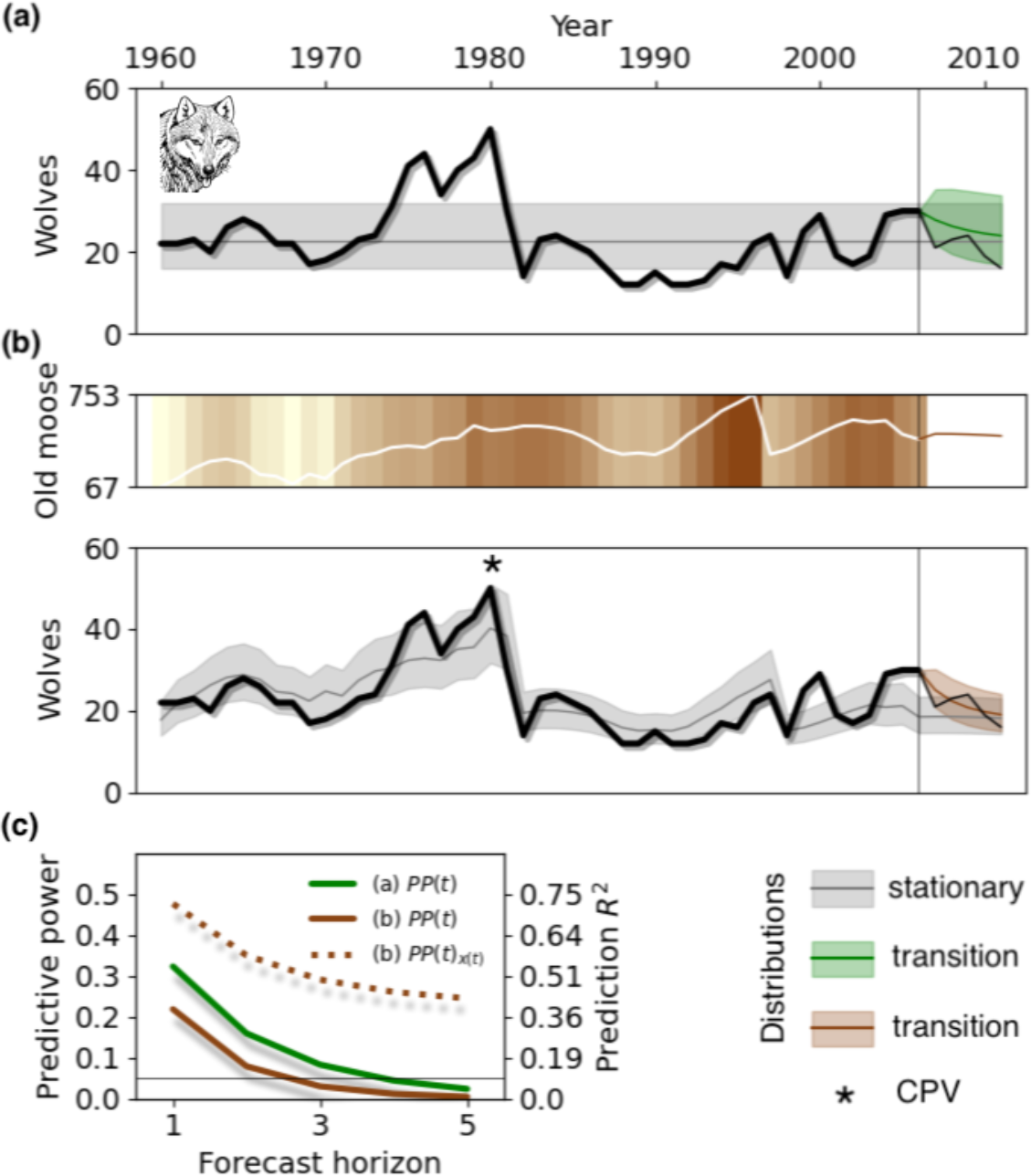
The main elements characterizing intrinsic predictability. Panel **(a)** depicts the abundance time series of the wolf population on Isle Royale (black line, 1959-2011), along with the stationary and transition distribution (legend bottom-right of figure). For illustrative purposes, we take the year 2006 as the ‘current’ year to make predictions (vertical line). Panel **(b)** depicts the analogous situation as in panel (a), except that now two covariates – moose population (top of panel) and effects of CPV (asterisk) – are also used in the computation. The transition distribution here is therefore conditional on the inclusion of covariates. Panel **(c)** gives predictive power values, *PP*(*t*), for the panels (a-b), and for panel (b) also *PP*(*t*)*x*(*t*) (derived in Box 1). All distribution variances in (a-b) are visualized as 66% confidence interval. Drawing: author unknown, public domain.

Many textbook deterministic models show how density-regulated populations eventually reach and stay at a ‘carrying capacity’. The stochastic analogue is the stationary distribution, with a mean (the ‘carrying capacity’) and a variance capturing, for example, yearly fluctuations in abundance around the mean (Fig. 1a). If the properties of a stochastic process do not change over time, the stationary distribution offers the simplest way of forecasting future abundances: just predict that the mean and uncertainty in the abundance of a species in any future year is given by the stationary distribution, regardless of the current abundance. This approach is easy to implement because the estimated sample mean and variance (and higher statistical moments) of the time series suffice to characterize the stationary distribution. A different but equally simple approach is to assume that next year’s abundance will be equal to the current one. Both of these approaches, however, are unlikely to be very accurate. Predictability, in a nutshell, takes a position between these two extremes by quantifying the predictive information available in a population time series.

When conditioning forecasts on current abundance, the forecast dynamics are characterized by the transition distribution, with mean and variance (also called forecast error variance: [31]) that change with the forecast horizon (Fig. 1a). Starting at the current abundance, the population is expected to eventually reach the stationary mean, and in the long run the transition distribution will completely overlap with the stationary distribution. These two distributions – transition and stationary – are the main elements characterizing predictability: the less the transition distribution overlaps with the stationary one, the more predictive information is available for reliable forecasts, and predictability is high; conversely, predictability is said to be lost when the transition distribution largely overlaps with the stationary distribution [32].

The predictability measure we use is the metric predictive power, *PP*(*t*), rooted in information theory and developed in climatology [31]. Previous work on predictability within an ecological setting has used model-free methods based on information theory [30,33] or computational irreducibility [29], methods that typically cannot be integrated into further modeling and forecasting for biodiversity conservation. An advantage of *PP*(*t*) is that we can associate predictability with specific dynamical patterns observed in a time series by computing *PP*(*t*) in terms of estimated model parameters (Box 1). Predictive power, 0 < *PP(t)* < 1, depends on the forecast horizon *t*, and *PP (t)* will decrease to zero as *t* increases to infinity and the transition distribution converges to the stationary distribution (Fig. 1c). In practice, it makes sense to define a predictability barrier that gives a threshold below which the predictability of the system is negligible. To cast predictability in familiar statistical terms, *PP (t)* can be shown to be related to the theoretically maximum possible prediction *R*^2^ reflecting forecast accuracy (Box 1). To determine a predictability barrier, we will use *PP (t)* = 0.05 as a threshold, corresponding to a maximum possible prediction *R*^2^ of ∼10% for univariate time series (Fig 1c).

We implement *PP*(*t*) using time-series models from the ARMA family [38] (Box 1). ARMA models are familiar to many ecologists and evolutionary biologists (e.g. [39]), and are used extensively due to their parsimonious and flexible nature. ARMA models make it possible to address many central topics in wildlife data, ranging from measurement error to population decline due to human stressors. For highly nonlinear processes like systems with multiple stable states (however elusive these are: [40]), ARMA models will fail to capture patterns caused by the nonlinearities and therefore may underestimate the true predictability of the data. This might argue for an approach that computes the information content of a dataset without the need to specify a model [30,33]. The cost of taking a model-free approach, however, is that there is no way to *directly* translate these information-based metrics into terms that assess model fit and predictive ability. Although we do not pursue this here, it is straightforward to compute *PP*(*t*) for nonlinear models where its interpretation and use is similar to ARMA models.

#### Box 1

**Predictive power and time series models**

The predictability metric predictive power, *PP*(*t*), has been developed using information-theoretic concepts to measure the uncertainty in predictions [31]. One advantage of this metric is that it can be easily reformulated for Gaussian processes like widely used time-series models (see below). For a univariate time series, the general multivariate Gaussian case (ESM Appendix 1.1, eq. S1) reduces to

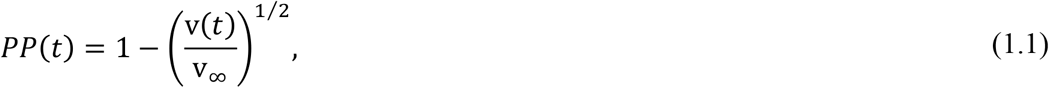

where v(*t*) and v_∞_ are the variances of the transition and stationary distributions, and *t* is the forecast horizon. This formulation focuses on variances and therefore implicitly assumes that the estimates of the distributions’ means are unbiased; see [31] for the inclusion of estimation uncertainty in *PP*(*t*). Calculation of v(*t*) and v_∞_ is outlined in the ESM Appendix 1.1, including simple expressions for *PP*(*t*) = 1. Where necessary, we will refer to *PP*(*t*) as the *intrinsic* predictability to distinguish it from other predictability metrics. For univariate processes, *PP*(*t*) can be related to the theoretical limit of forecast accuracy, *R*_*pred*_^2^(t) = 1− v(*t*) v_∞_ −1 [41], as *R*_*pred*_ ^2^(*t*) =1− (1 − *pp*(*t*))^2^. More generally, *PP (t)* declines over time (i.e. forecast horizon), eventually approaching zero: in this limiting case both distributions completely overlap and predictability is lost (Fig. 1). Alternatively, a loss of predictability could be defined using a threshold value set by *R*_*pred*_^2^(t), where for example a value of 0.10 would be considered a poor prediction. These findings – the link between *PP (t)* and forecast accuracy as measured by *R*_*pred*_^2^(t), and the decrease of predictability over time – theoretically confirm previously proposed but unresolved hypotheses ([36], and references therein).

Because eq. 1.1 is suitable for Gaussian processes, time-series models from the ARMA(*p,q*) family are apt candidates for implementing *PP* (*t*), and these linear models can also adequately represent nonlinear processes [38]. For the purpose of assessing predictability, it is convenient to state an ARMA(*p,q*) process as a dynamic regression model,

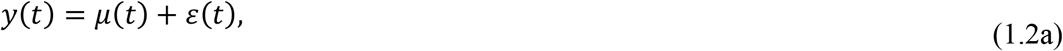

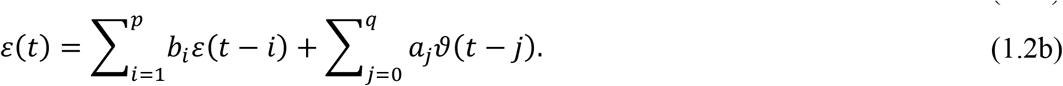

Here, *μ*(*t*) is the potentially time-varying mean of the process *y*(*t*) (Box 3), and *ε t* are temporally autocorrelated errors; without a dependence of the mean on time, *μ*(*t*) is simply a constant (e.g. ‘carrying capacity’). In cases where explanatory variables are available (section §2.2), eq. 1.2a becomes

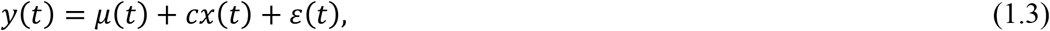

where *x(t)* is a covariate time series and *c* its coefficient. For this case, we can define a covariate predictive power

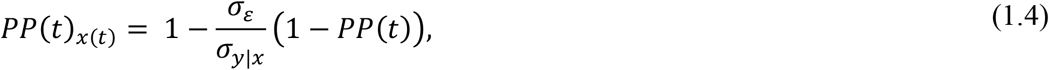

where *σ*_ε_ is the standard deviation of the random errors of the fitted model to the observed *y*(*t*) (eq. 1.2b), and *σ*_*y*|*x*_ is the standard deviation of the overall stationary distribution of *y t* that depends on variation in *x*(*t*) (ESM Appendix 1.2).

For a univariate ARMA model, *PP*(*t*) depends on the *p* + *q* ARMA coefficients (eq. 1.2b), whereas the residual variance of the process 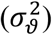 cancels out (ESM Appendix 1.1). From a simulation study presented in the ESM Appendix 1.4, we recommend setting *q* = 0 for unbiased estimates of predictive power.

### 2.2 Predictability when explanatory variables are available

Outside the context of time-series forecasting, predictability is often associated with ‘predictor’ (explanatory or independent) variables; for example, simple regression models ask how the mean of a response variable *y t* depends on an explanatory variable *x t*. In contrast, predictability as measured by *PP t* depends on the stochastic properties of a model (Box 1, eq. 1.2-1.3). To avoid confusion between ‘predictor’ variables and predictability, we need to discuss predictability in models with explanatory variables.

To make multi-year forecasts, an explanatory variable *x*(*t*) is only useful if we know its future values, ideally without error. Without any information about the future values of *x*(*t*), even if it explains a lot of variation in the observed values of *y*(*t*), it provides no information for forecasting. Therefore, forecasting can be performed using a model fit to the observed values of *y*(*t*) that ignores *x*(*t*) altogether, and the issue of explanatory variables evaporates. Consider, though, the opposite extreme in which we know the future values of *x*(*t*) precisely. While this might sound unlikely, it will be the case if we are making forecasts under different scenarios, in which case *x*(*t*) might be, for example, given in a scenario of future climate change. Alternatively, future ‘known’ values can come from a deterministic projection of *x t* into the future given the dynamics of *x*(*t*) so far (Fig. 1b).

When incorporating an explanatory variable, care must be taken in interpreting predictability. For example, *x*(*t*) might explain a substantial part of the patterns in the observed values of *y*(*t*) that would otherwise be attributed to the random errors (Box 1, eq. 1.2a-b). If these patterns generate the predictability captured by *PP*(*t*), incorporating *x*(*t*) could simultaneously increase the overall fit of the model to the observed *y*(*t*) and decrease *PP*(*t*) (Fig. 1b,c). However, this seeming incongruence only occurs because it ignores the component of the forecast that uses the future values of *x*(*t*) to predict changes in the mean of the stationary distribution. In Box 1 (eq. 1.4) we present an extended version of predictive power, the covariate predictive power *PP*(*t*)_x(t)_, which captures the overall predictive power in the presence of *x*(*t*).

### 2.3 Univariate or multivariate time series?

In certain situations multiple correlated time series may be available, for example for an ecological community or for different phenotypic measures of an organism. How should predictability be assessed for such situations? Should all time series be analyzed separately? Should the dimensionality be reduced using a principle component analysis (PCA) or similar approach, and subsequently only the first component used? Or should all data be concomitantly analyzed using a multivariate ARMA(*p,q*) model (ESM Appendix 1.1)?

As an illustrative example we analyzed three beak measures of three Galapagos finch populations on Daphne Major Island for the period 1973-2012 [42] (ESM Fig. S2, Appendix 4). Analyses of the medium ground finch (*Geospiza fortis*) and small cactus finch (*G. scandens*) give similar results for all three approaches (Fig. 2): analyzing traits separately, first performing a PCA and using the first component, and analyzing all three traits together. For the large ground finch (*G. magnirostris*), however, the multivariate assessment clearly leads to greater predictability, because there is information in the co-variances of the traits through time.

**Fig. 2.**
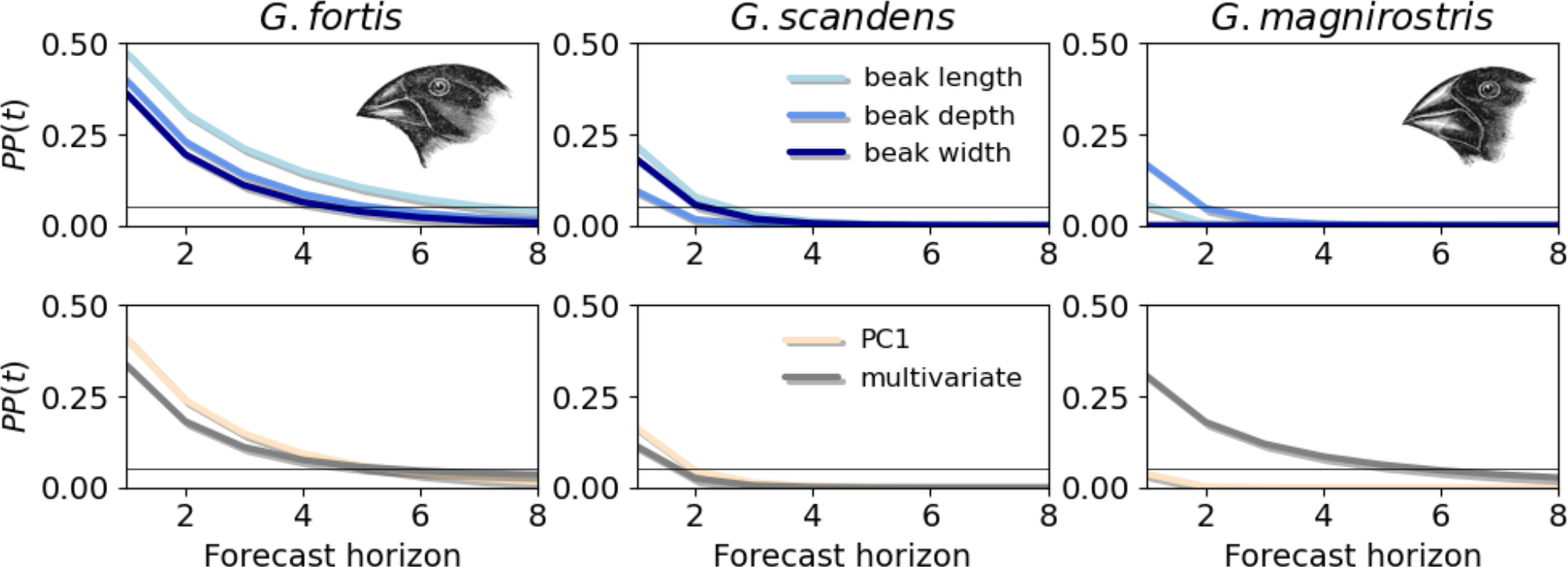
Predictability of univariate vs. multivariate time series. The first row gives predictability results for each species (column title) with all three measures analyzed separately, and the second row gives analogous results based on the first component of a PCA (PC1) and the multivariate assessment. Drawings: John Gould, public domain.

This example illustrates that, while there is no guarantee that a multivariate approach for individual trait or ecological community data provides higher predictability, it would be prudent to try. Further, note that predictability of multivariate datasets as measured by *PP t* also allows distinguishing predictable and unpredictable components of the system by performing a predictable component analysis, reminiscent of a PCA [31,43]. For data on an ecological community, as an example, such approach would make it possible to parse out the leading drivers of community predictability.

## 3. Limit of predictability across animal taxa

To gain a bird’s eye view on predictability of animal population sizes and phenotypes, we analyzed 320 invertebrate and 963 vertebrate ecological time series from the four taxonomic groups insects, birds, mammals and fish, where invertebrate data were either collected at the species level or at the plot level [44,45] (ESM Fig. S3, Table S1, Appendix 3); and we analyzed 307 phenotypic time series of bird, fish, and mammal populations [46] (ESM Fig. S3, Table S3, Appendix 3).

For both ecological and phenotypic time series, we found that 50% of all analyzed populations are predictable at most one year ahead, with interquartile ranges of 0-3 years and 0-2 years, respectively (ESM Fig. S4). Because values of the predictability barrier and one-year-ahead predictive power, *PP*(1), are strongly associated (Spearman’s *ϱ* = 0.95, *P* < 0.0001), in Fig. 3 we present *PP*(1) values separated by taxonomic groups, and for ecological time series also by illustrative sub-groups/families.The results indicate that mammal populations have the highest predictability, followed by fishes, birds, and insects (ecological data only). All results combined (grey lines in Fig. 3), 50% of ecological time series have a *PP*(1) value of at most 0.13, and 0.06 for phenotypic time series. These are sobering results: *PP*(1) = 0.1 translates to a maximum possible prediction *R*^2^ of ∼20% (Box 1), for the humble endeavor of predicting next year’s ecological or phenotypic population value.

**Fig. 3.**
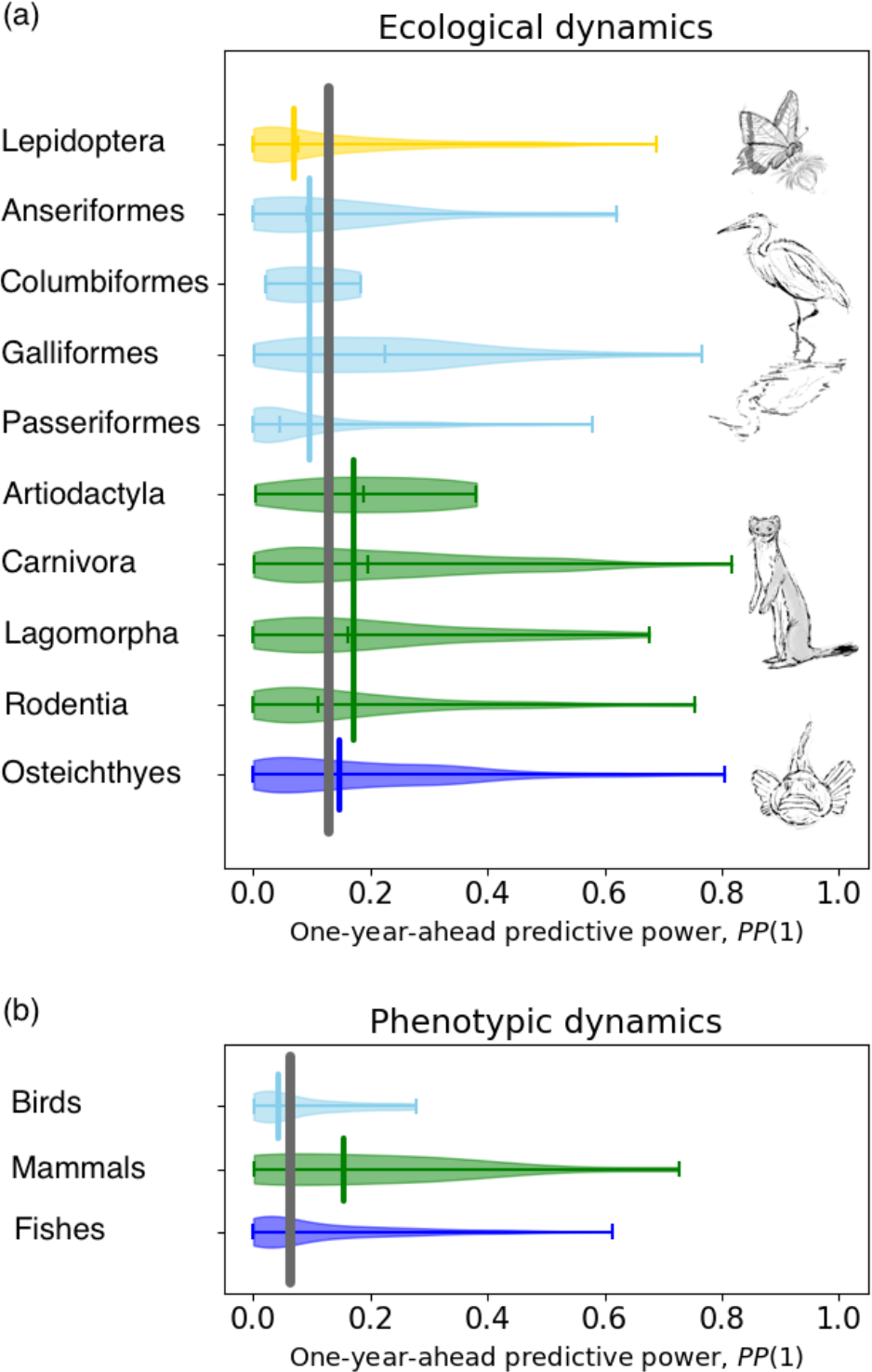
Predictability across animal populations. The distributions of one-year-ahead predictive power values, *PP*(1), are shown for (**a**) ecological and (**b**) phenotypic population time series, sorted by the four taxonomic groups insects (only panel a), birds, mammals and fish, and in panel (a) additionally sorted by 10 illustrative taxonomic subgroups. For each taxonomic group a vertical line gives the median value over all respective populations; smaller vertical lines give the median and min/max values. In both panels, the overall median value is marked with a vertical grey line. Drawings: Andrea Klaiber, **©** Wildlife Analysis GmbH, Switzerland.

To better understand the variation in *PP* 1 illustrated in Fig. 3, we inspected its association with ecological characteristics; for illustrative purposes we focus on ecological dynamics. Primary threats (human stressors) and the total number of threats for birds, fishes, and mammals ([4]; ESM Table S2) lacked evident patterns with predictability (ESM Fig. S5). On the other hand, freshwater populations of all taxonomic groups are characterized by lower predictability compared to other realms (Fig. 4a); nonetheless, this could be driven by the proportional dominance of insects in the sample (Fig. 3, ESM Table S1). For insect populations, we combined realm information with protection status (Fig. 4b) or with data level (Fig. 4c; single populations vs. dynamics of all insects in a monitored plot): in both cases, freshwater species again appear to be less predictable. Furthermore, while terrestrial species seem to be more predictable in protected areas, the data level does not seem to influence predictability, the latter result being good news for insect monitoring which often relies on plot-level data. This result, however, does not support a previously proposed hypothesis that aggregated data may be more predictable ([36], and references therein). Finally, populations along the Pacific coast in the northern hemisphere appear to be less predictable than populations in the North Atlantic (Fig. 4d), and this pattern is driven by fish populations (ESM Fig. S6, Appendix 3): our results suggest that along the Pacific coast population dynamics harbor less effects of the food webs in which they are embedded as opposed to the North Atlantic. This finding offers statistical evidence for a previously proposed hypothesis that the food web embedding may affect predictability ([36], and references therein).

**Fig. 4.**
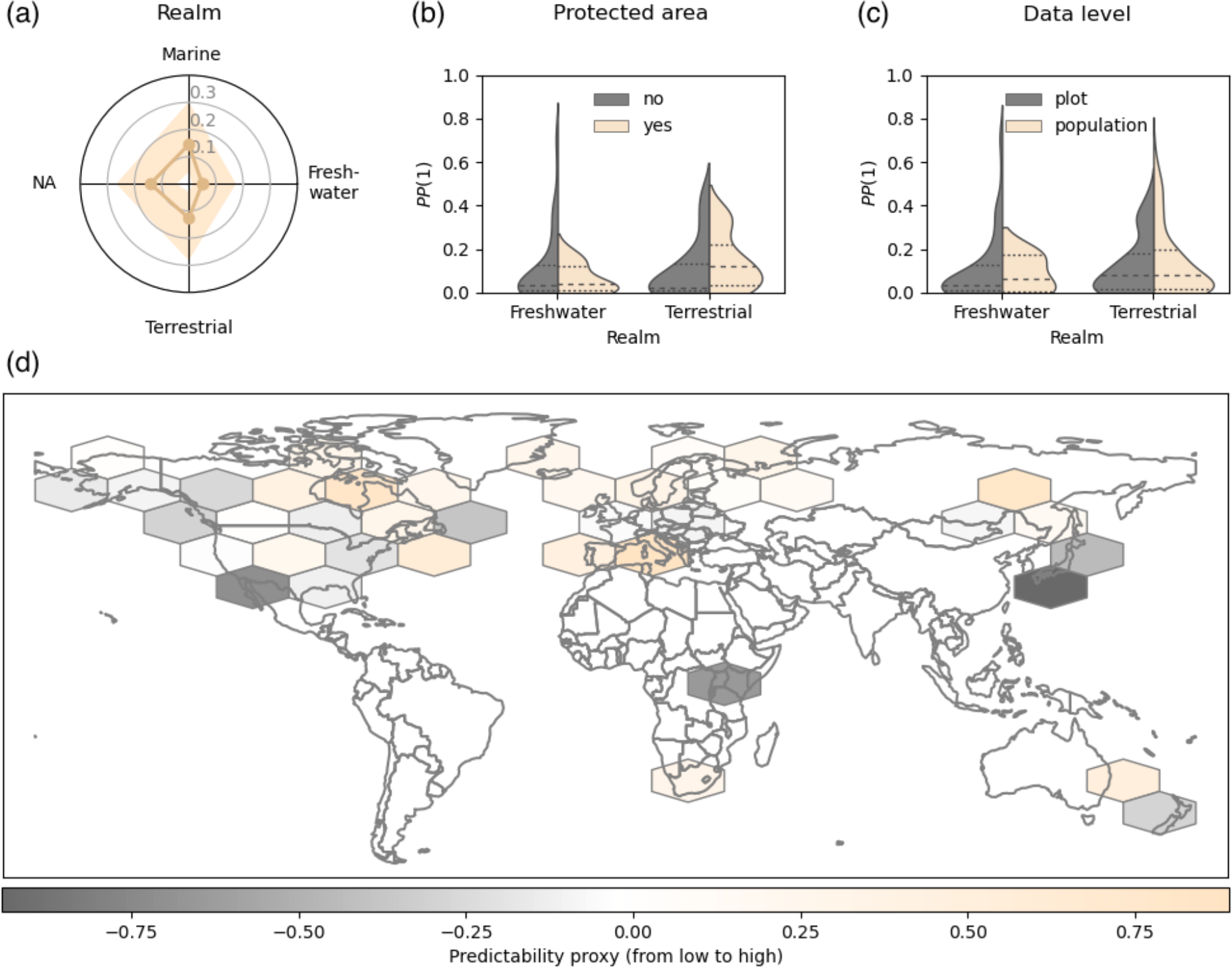
Inspecting the predictability of ecological dynamics. The radial graph (**a**) summarizes the one-year-ahead predictive power values, *PP*(1), sorted by realm (median and interquartile range). The two split violin plots (**b-c**) depict the distribution of *PP*(1) values of insects, sorted by protection status and data level (species or plot time series), respectively (median and interquartile range), in both cases further classified by realm. The world map (**d**) depicts, as a hexagonal grid, a transformation and summary of geographically sorted *PP*(1) values (see color bar).

## 4. Exploited populations

Billions of people worldwide rely on and benefit from the use of about 50,000 wild species for multiple purposes including food, medicine, and recreation [47]. Unfortunately, exploitation has become a main cause of elevated extinction risk, affecting species from many faunal and floral taxonomic groups (e.g. [4,48,49]). For example, while well-assessed fisheries are moving towards sustainability, 80% of the global fish catch is derived from non-assessed populations of which especially small fisheries are in a worrisome state [50].

Wildlife management has increasingly sought harvesting strategies to minimize overexploitation, and nowadays many approaches include forecasting population dynamics to abide by sustainable management (e.g. [24,51,52]). We expect predictability of exploited populations to reflect life history characteristics: because larger animals tend to be exploited relatively more often [53] and their life history characteristics correlate with a decreased degree of density dependence [54], this will lead to higher predictability (Box 1). But could it be that different harvesting strategies affect predictability to varying degrees (Box 2)?

### Box 2

**Harvesting strategies and predictability**

One way of categorizing harvesting strategies is to relate annual harvest targets to the (estimated) population abundance. For example, in a constant yield strategy a fixed number of animals is harvested irrespective of abundance, maybe with the exception of stopping outtakes if the population is estimated to undershoot a critical low level (threshold). A different strategy is to harvest a fixed proportion of animals, where the proportion does not depend on the population size, but the realized harvest (hunting bag) will; here again, the outtakes may be stopped as described above. As we explain below, different harvesting strategies can affect predictability differentially.

It is convenient to express harvest, *H*(*t*), as the product of the proportion of the population harvested, *p*(*t, N*(*t*)), and population size, *N*(*t*): *H*(*t*) = *p*(*t, N*(*t*)) *N*(*t*), where *p*(·) is potentially a function of *N*(*t*). Here, we consider the case in which *p*(*t,N*(*t*)) can be approximately expressed as a linear function of *y*(*t*) = ln (*N*(*t*), so that *H*(*t*) = h_0_ + *h*_1_*y*(*t*) + *𝒽* (*t*))*N*(*t*) and *𝒽*(*t*) are the residuals of the fitted proportion. If *h*_1_ = 0, we recover the abundance-independent proportional strategy. For *h*_1_ > 0, the proportion harvested increases with increasing *N*(*t*), and the intercept *h*_1_ sets the threshold below which no animal is harvested. Finally, *h*_1_ < 0 gives a linear approximation to a constant yield strategy at equilibrium, *p*(*t,N*(*t*)) = *Q*(*t*)*N*(*t*)^-1^, where *Q*(*t*) is the constant yield. This model for a harvested population generates the autoregressive parameter *b*_1_ which depends on the harvesting parameter *h*_1_, *b*_1_ =(1 − *h*_1_*e*^-β^) (1 – *γ*), where *β* is the birth rate and (1 – *γ*) is the autoregressive parameter of the unharvested population (ESM Appendix 5.1). Because parameter *b*_1_ directly affects predictability (Box 1), harvesting strategies with *h*_1_ ≠ 0 will do the same.

To investigate the potential effect of harvesting strategies on predictability, we analyzed 14 Swiss cantonal populations of the northern (Alpine) chamois (*Rupicapra rupicapra*), for which hunting bag and abundance time series are available [55]. To determine how harvesting strategies affect predictability, we computed predictive power for a population twice, with and without harvesting (ESM Appendix 5.2). The geographic distribution of one-year-ahead predictive power values, *PP*(1), reveals no intuitive pattern (Fig. 5a). The harvesting strategy captured by the harvesting parameter *h*_1_ (Box 2), however, evidences a clear effect on predictability (Fig. 5b): for populations subject to a harvesting strategy in which the proportion harvested is precautionarily adjusted according to the current abundance (*h*_1_ > 0), predictability is decreased, while populations subject to constant yield-like harvesting (*h*_1_ < 0) have increased predictability. This result occurs because the state-dependent management approach tracks the yearly population dynamics to decrease the variation in population fluctuations which thus decreases predictability (Box 1). In contrast, a constant yield-like harvesting strategy does not dampen population fluctuations but instead increases them relative to the mean population by depressing the population, thereby increasing predictability. As with all harvested species, a state-dependent harvesting strategies is more likely to lead to sustainability and reduced extinction risk (e.g. [51]). Thus, the loss of predictability caused by state-dependent management is an indication that management is effective. Therefore, predictability should not be the goal of management, but sustainable practice must be.

**Fig. 5.**
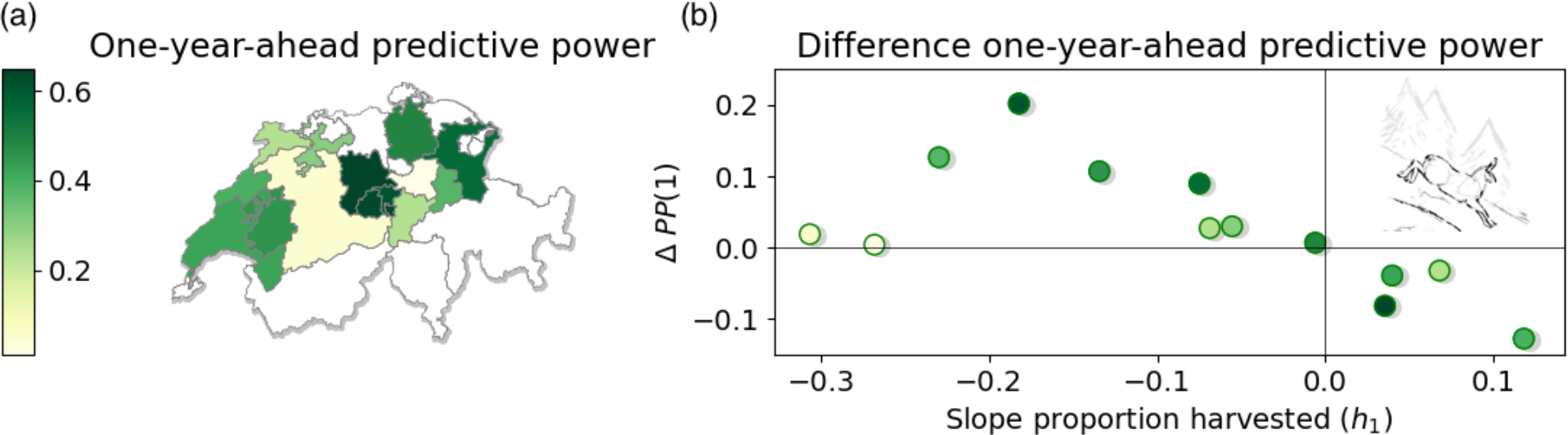
Effect of harvesting strategy on predictability. Panel **(a)** geographically presents estimated one-year-ahead predictive power values, *PP*(1), of 14 Swiss cantonal chamois populations. Panel **(b)** illustrates the change in the one-year-ahead predictive power, Δ*PP*(1) (harvested minus non-harvested), as a function of the harvesting strategy characterized by the slope value (*h*_1_) on the *x*-axis (i.e. how the proportion harvested depends on abundance; Box 2). The dot colors are the same as in panel (a). Drawing: Andrea Klaiber, **©** Wildlife Analysis GmbH, Switzerland.

## 5. Predictability of forced systems

So far, we have treated a potentially time-varying mean as a ‘nuisance’, which must be accounted for to compute the intrinsic predictability of a process (Box 1). In terms of assessing the predictability of the future dynamics, we implicitly assumed that the projected mean would be known. In a biodiversity conservation context the long-term trend of a population (abundance or phenotype) – a so-called forced system – is of central importance [39]. Time trends can for example be driven by habitat degradation, by inbreeding depression, or by introduced generalist predators [56–58]. Our confidence in projected mean changes, however, will be affected by uncertainty, but this aspect has been largely ignored in the ecological predictability literature (e.g. [30,36]). Compared to unforced systems (Fig. 1), for the predictability assessment of a forced system we now have a transition distribution that approaches the *forced* stationary distribution with time, and in addition this forced stationary distribution will separate from the current stationary distribution (Fig. 6a,b). These three distributions allow computing two complementary predictability components: (i) as before, the intrinsic predictability assesses our predictive ability *around* the (changing) mean, while (ii) the forced predictability assesses our ability to infer a stationary distribution with a changing mean (Box 3). If the dynamics *around* the mean are unpredictable, then the only way of predicting future population values is by using the forced mean.

**Fig. 6.**
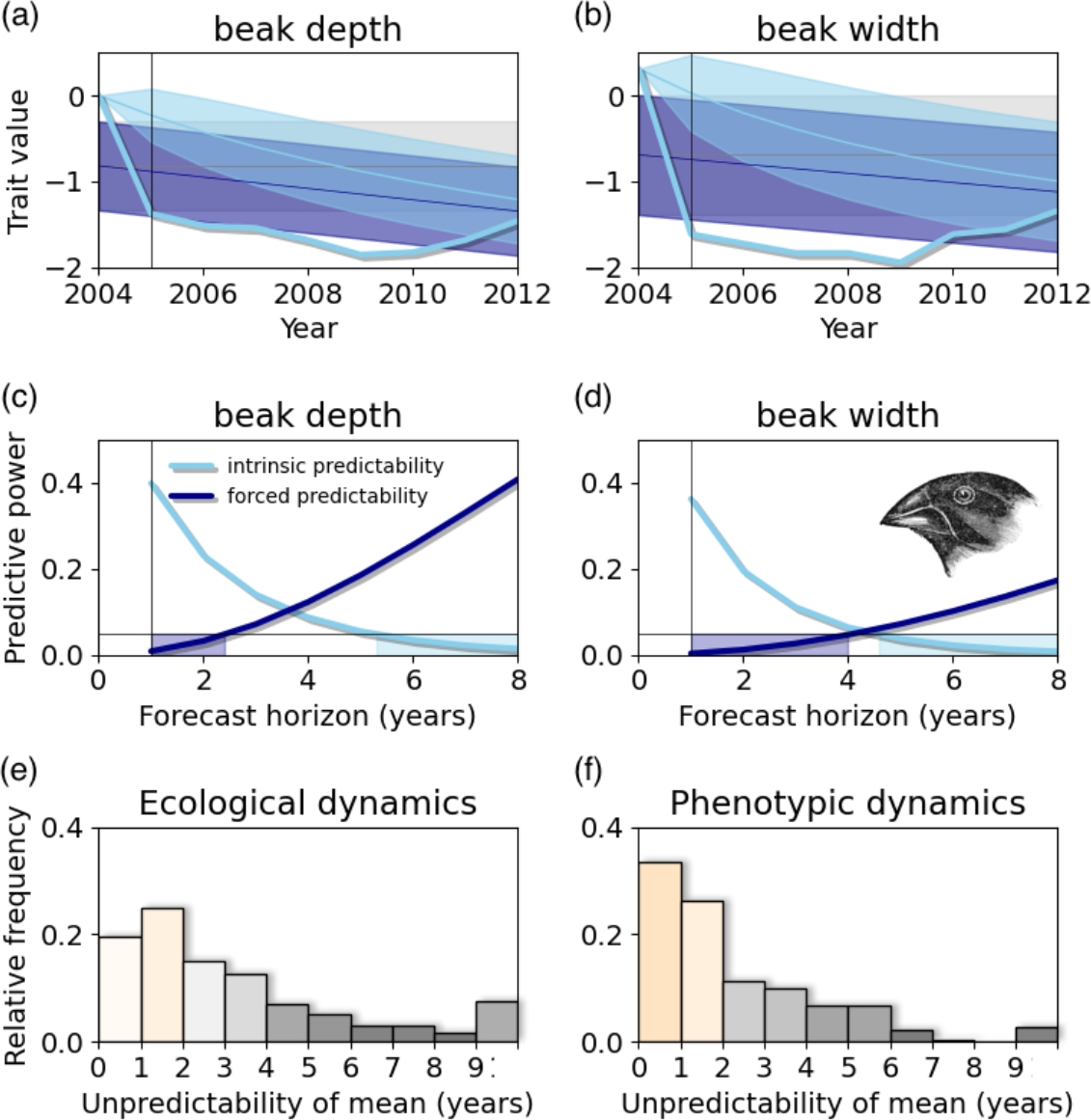
Predictability of forced systems. For two beak measures (panel title) of the medium ground finch (*G. fortis*), panels (**a-b**) depict the z-transformed data (2004-2012; ESM Fig. S2), the current stationary distribution (in grey), the forced stationary distribution (dark blue), and the transition distribution (light blue). All distribution variances are visualized as 66% confidence interval. Panels (**c-d**) give the intrinsic predictability, *PP t*, and the forced predictability, 1 − exp −*M*_F_ (legend; Box 3), for the two beak measures. The colored areas correspond to the unpredictable period of the forced (dark blue) and intrinsic (light blue) predictability, respectively. In panels (a-d), the current year is assumed 2004, from which forecast would be made. Panels (**e-f**) show the duration of the initial unpredictable period of the forced predictability (see the dark blue areas in panels c-d) for all ecological and phenotypic time series (title) with a time trend. Drawing: John Gould, public domain.

### Box 3

**Processes with a time-varying mean (forced systems)**

Predictive power is defined for stationary processes, although the definition can be extended for processes with a time-varying mean such as a declining trend in population abundance (Box 1, eq. 1.1-1.2). A simple, robust phenomenological approach is to include a deterministic time trend *μ t* as a covariate into eq. 1.2a [59], where

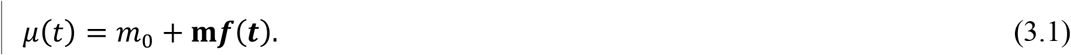

Here, for a linear time trend the second term on the right-hand side of eq. 3.1 becomes *m*_1_*t*, and for a quadratic (i.e. nonlinear) time trend this term becomes *m*_1_*t* + *m*_1_*t*^2^. For a forced system, predictive power can be extended to also account for the continued (forced) change in the mean, i.e. a forced stationary distribution [60]. This case is closely related to predictive power when there is an explanatory variable, *PP*(*t*)_x(*t*)_ (Box 1, eq. 1.4): for *PP*(*t*)_x(*t*)_ the covariate data *x*(*t*) enter the analysis as scalar values, whereas to compute forced predictability the whole forced stationary distribution is used, not just a changing scalar mean. The total predictability of a forced system as measured by predictive power is

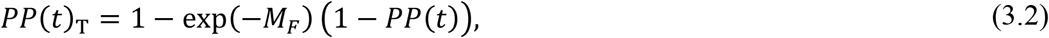

where *M*_F_=(*μ*(*t*) – *μ*(0))^2^ (2v_∞_)^-1^, *μ* 0 is the current mean,*μ t* si the projected mean *t* years ahead, and v_∞_ the variance of the stationary distribution (Box 1). Without a changing mean (*M*_F_ = 0), eq. 3.2 reduces to *PP*(*t*). Intuitively, our ability to predict a forced mean increases with more pronounced mean changes and decreases with more variable dynamics. In addition to combining both aspects of predictability into one metric *PP*(*t*)_T_, it is also informative to compare both components of *PP*(*t*)_T_, that is, comparing *PP*(*t*)_T_ = *PP*(*t*) when *M*_F_ = 0 to *PP*(*t*)_T_ = 1 − exp −*M*_F_ when *PP*(*t*) = 0. These two components reveal opposite behavior: while *PP*(*t*) decreases to zero with time, the forced predictability increases continually away from zero if the forcing persists (Fig. 6). Therefore, contrasting both measures reveals what aspect of the dynamics is worth focusing on for predictions (ESM Appendix 1.3).

To illustrate the predictability of forced systems, we start with the three beak measures of the medium ground finch (*G. fortis*; section §2.3), all analyzed separately. While beak length statistically lacks a time trend, both beak depth and width decrease over time (ESM Fig. S2). For the latter two beak measures, panels (a-b) in Fig. 6 depict the respective three distributions used for assessing predictability, and panels (c-d) give the intrinsic and the forced predictability separately. For beak depth, changes in the mean are largely unpredictable for 2 years (i.e. the forced predictability value is below the threshold value of 0.05; Box 1), while for beak width they are unpredictable for 4 years; the intrinsic predictability *PP t* in both cases lasts ∼5 years. Confronting the two predictability metrics shows that, to predict these two phenotypes more than 4 years into the future, using the forced mean gives more information than the current trait values.

Because many biodiversity assessments focus on long-term trends [17,39], and because many populations show a lack of intrinsic predictability (Fig. 3), a simple way of using forced predictability is to just ask about the number of forthcoming years (or generations: ESM Fig. S8) during which a changing mean is unpredictable (Fig. 6c,d). To investigate this question broadly, we again used the data collections from section §3 selecting time series with statistically detectable trends (*n* = 948). We found that approximately half of the populations had an unpredictable mean change for at least one year, and potentially many more (Fig. 6e,f); predicting a trend 1-2 years ahead might therefore be futile in many cases.

## 6. Why predictability matters for biodiversity conservation

For any biodiversity change assessment, the first step should be to evaluate predictability. Predictive power, *PP*(*t*), gives an objective summary of the information content in the data and a first cut at more-detailed modeling to make predictions. *PP*(*t*) focuses on the ‘unexplained’ variation in a time series that is not associated with any explanatory variable, asking whether this variation can be used for predictions (Box 1). *PP*(*t*) can also be used with models including explanatory variables (section §2.2) or forcing terms (section §5), to assess where the focus and further data-related efforts should be placed to make predictions. Thus, *PP*(*t*) can parse out where informative patterns exist in time-series data, directing attention to how reliable and credible predictions can best be made.

The predictability measure *PP*(*t*) is generic, in the sense that it can be applied to any time series or simultaneously to multiple time series. Our finding that the predictability of the ecological and phenotypic data collections was similar in magnitude surprised us (Fig. 3), because we expected population-level phenotypic changes to be too slow to be detectable in most time-series datasets. However, if the fitness consequences of traits are large, then traits should change on similar time scales as population abundances [27]. Less surprising were the generally low *PP*(*t*) values (Fig. 3), confirming previous findings [29] but also reflecting the heterogeneity of quality among time series and their short lengths. This underscores the need to compute *PP*(*t*) in biodiversity conservation where time series of interest are often short and of suboptimal quality [15]. Low *PP* (*t*) values might simultaneously caution against further statistical analyses and argue for future concerted efforts to increase predictive information in the data; a ‘better model’ will be of little help for data that are inherently unpredictable.

Predictability, however, is not an immutable property of a population or phenotypic trait; instead, it depends on the forces that affect populations and on how much data are available. For example, for the case of exploited populations we have explained how the harvesting strategy can change predictability. We have also illustrated how considering explanatory variables and time trends (forcing terms) or performing a multivariate analysis may increase predictability. Although data for explanatory variables or multiple time series might be a rare luxury in biodiversity conservation, intrinsic and forced predictability can provide a robust means to assessing the limit of predictive ability.

What are some fruitful ways to put predictability measures into practice? At the level of whole data collections, sensible versions of *PP*(*t*) (Box 1, Box 3) will be useful as population-level weights to compute synoptic indicators of projected biodiversity changes (e.g. [17]), to appropriately ‘flag’ population data with low predictive information and, therefore, to denote the credibility of the projections. As a second example, the distribution of *PP*(*t*) values among taxonomic groups (Fig. 3) indicates that it may be worth adopting group-specific predictive approaches. For example, many insect populations show low intrinsic predictability, suggesting focus should be placed on forced predictability (i.e. predictions of a changing mean); it may especially be worthwhile exploring ways of considering explanatory environmental variables like habitat or climate characteristics to increase predictive ability. Other species like mammals, on the other hand, may more frequently warrant employing predictions conditioned on the present population value (intrinsic predictability), in addition to or in combination with predicting long-term trends (Box 3).

In conclusion, if predictability metrics are not adopted, biodiversity conservation risks turning the horsemen of the environmental apocalypse into ghost horsemen that might be invisible or be a mirage, mistakenly seen even when they are not there. As scientists worried about biodiversity loss, we often call for more and higher-quality data; what we really want is data that contain more predictive information. Given the urgency for action dictated by a rapidly changing world, we need to use the hard-won data we already have and explore ways of adding predictive information to these data – we cannot afford to wait for ‘better’ data to become available while human-caused stressors are wreaking havoc on wildlife. We believe that embracing the concept of predictability will benefit the urgent and global endeavor of protecting life on Earth.

## Supporting information

ESM

## Data availability

The data that support the findings of this study are publicly available online and are referenced in the main text and the ESM Appendix 6.

## Code availability

Code to compute predictability metrics (in Python and in R) will be uploaded to the zenodo repository after acceptance.

## Acknowledgments

We thank Thorsten Hens for valuable suggestions on improving the manuscript.

## Author contributions

**C.B**. Conceptualization, Methodology, Formal analysis, Data Curation, Writing - Original Draft, Writing - Review & Editing, Visualization. **A.R.I**. Methodology, Formal analysis, Writing - Review & Editing.

