## Supplementary material for "Predictability of ecological and evolutionary dynamics in a changing world": ESM

**Electronic supporting material for**

**This PDF file contains:**

- Supplementary Tables S1-S3
- Supplementary Figures S1-S8
- Appendix 1: Computation of predictive power
  1.1 Without explanatory variables
  1.2 With explanatory variables
  1.3 Predictability of forced systems
  1.4 Simulation study: AR(*p*) vs. ARMA(*p*,*q*)
- Appendix 2: Analysis of wolf population on Isle Royale
- Appendix 3: Analysis of Galápagos finch phenotypic time series
- Appendix 4: Analysis of ecological and phenotypic time series collections
- Appendix 5: Exploited populations
  5.1 Dynamical model of an exploited population
  5.2 Analysis of Alpine chamois populations
- Appendix 6: Data sources
- References electronic supporting material

#### Supplementary tables

**Table S1. Ecological time series categorized by four aspects.** All time series analyzed in this study are here ordered by (i) taxonomic group (rows), (ii) insects additionally by data level (species-level or plot-level; two separate rows), (iii) realm (first group of columns), and (iv) protection status of the habitat (insects only; second group of columns). 'NA': no information available; ' – ': no time series available for a particular category. Original data sources: see Appendix 6.

| | Realm | | | | Protected area | | |
|---|---|---|---|---|---|---|---|
| | Terrestrial | Freshwater | Marine | NA | Yes | No | NA |
| **Insects** | 211 | 76 | – | 33 | 34 | 88 | 198 |
| species | 160 | 5 | – | 33 | – | – | 198 |
| plot | 51 | 71 | – | – | 34 | 88 | – |
| **Birds** | 199 | 5 | 12 | 37 | – | – | – |
| **Mammals** | 401 | 3 | 8 | 71 | – | – | – |
| **Fish** | – | 11 | 216 | – | – | – | – |

**Table S2. Populations and threats**. For a given primary threat or the number of co-acting threats (columns), the table shows the proportions of the three taxonomic groups (rows) within each column; the columns sum to one (except for rounding differences). All proportions have been computed using the data published in [1].

| | Primary threat | | | | | Number of threats | | | |
|---|---|---|---|---|---|---|---|---|---|
| | Climate | Exploitation | Habitat | Invasion | Pollution | 0 | 1 | 2 | 3 |
| **Birds** | 0.611 | 0.123 | 0.543 | 0.455 | 0.751 | 0.383 | 0.296 | 0.529 | 0.495 |
| **Mammals** | 0.082 | 0.310 | 0.278 | 0.421 | 0.166 | 0.332 | 0.260 | 0.272 | 0.370 |
| **Fish** | 0.307 | 0.567 | 0.178 | 0.124 | 0.083 | 0.285 | 0.444 | 0.199 | 0.135 |

**Table S3. Phenotypic traits**. For the three taxonomic groups birds, mammals, and fish (Fig. S3) the number of analyzed phenotypic time series is shown by trait type; for further details on these trait type categories, see [2].

| | **Phenology** | **Behavior** | **Physiology** | **Growth** | **Size** | **Other morphology** | **Other life history traits** |
|---|---|---|---|---|---|---|---|
| **Birds** | 13 | – | – | – | 9 | 22 | 8 |
| **Mammals** | – | – | – | – | 7 | 9 | – |
| **Fish** | – | – | – | 34 | 166 | 11 | 28 |

#### Supplementary figures

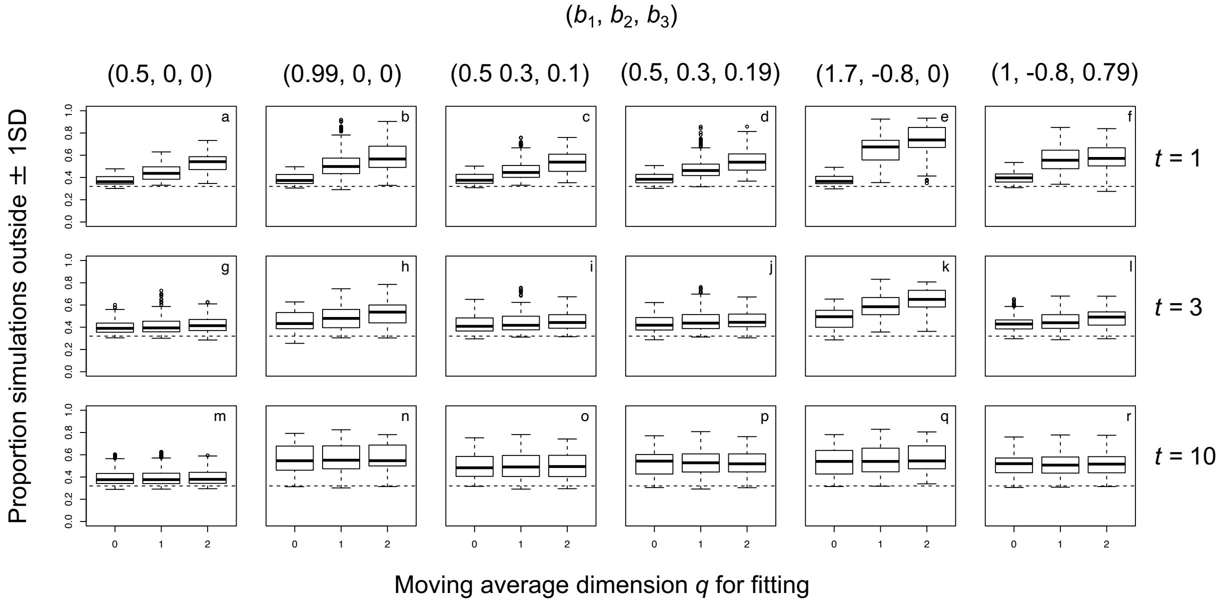


**Fig. S1. Estimating predictive power with AR(*p*) vs ARMA(*p*,*q*).** We simulated datasets using ARMA(*p*,*q*) models for six sets of AR coefficients ($b_1$, $b_2$, $b_3$; column titles). These six sets are made up of three pairs, with the second of each pair having a stationary distribution with greater variance. The sets of AR coefficients were crossed with nine sets of MA coefficients to give 54 parameter combinations that generate distinct dynamics. For each of the 54 parameter combinations, we included a linear time trend by letting $\mu(t) = m_0 + m_1 t$, where the coefficient $m_1$ took values 0, –2.5, –5.0, –7.5, and –10.0. Finally, we considered time series of length 30 and 100, for a total of 540 scenarios, and 1000 simulations were performed for each scenario. To assess the accuracy of the estimate of the transition distribution, for each simulated time series we scored whether the actual value fell within one standard deviation of the mean of the estimated transition distribution at forecast horizons of $t$ = 1, 3, or 10. If the estimate of the transition distribution is accurate, then 33% of the projected values of the time series should fall outside one standard deviation of the mean of the transition distribution. The histograms show the results for all 540 scenarios fitted with ARMA(*p*,*q*) models having $q$ = 0, 1, or 2, with separate panels per row aggregating the simulations into the six groups with different AR coefficients, and rows aggregating results per forecast horizons of $t$ = 1, 3, or 10.

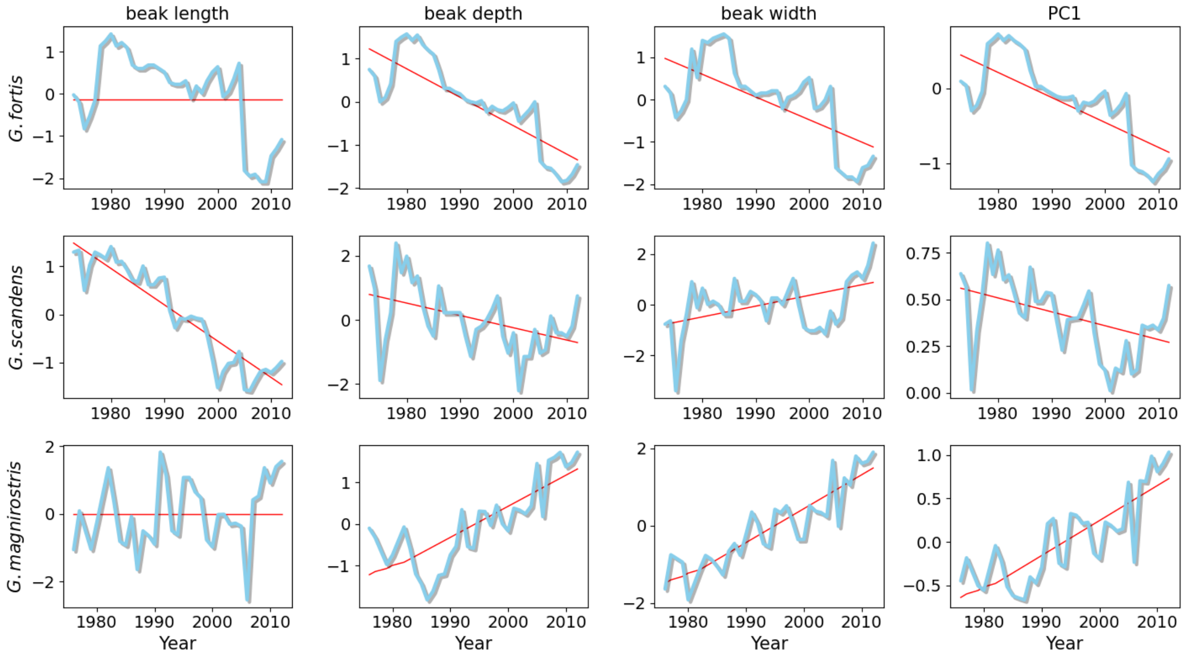


**Fig. S2. Phenotypic time series of three Galápagos finch species on Daphne Major Island.** The first three columns show three z-transformed mean beak trait time series (column labels) of three Galápagos finch species (row labels), measured on Daphne Major Island (1973-2012) by Peter and Rosemary Grant [3]. The last column shows the first component of a principal component analysis (PC1) of the three beak traits, also from [3]. Data source: see Appendix 6.

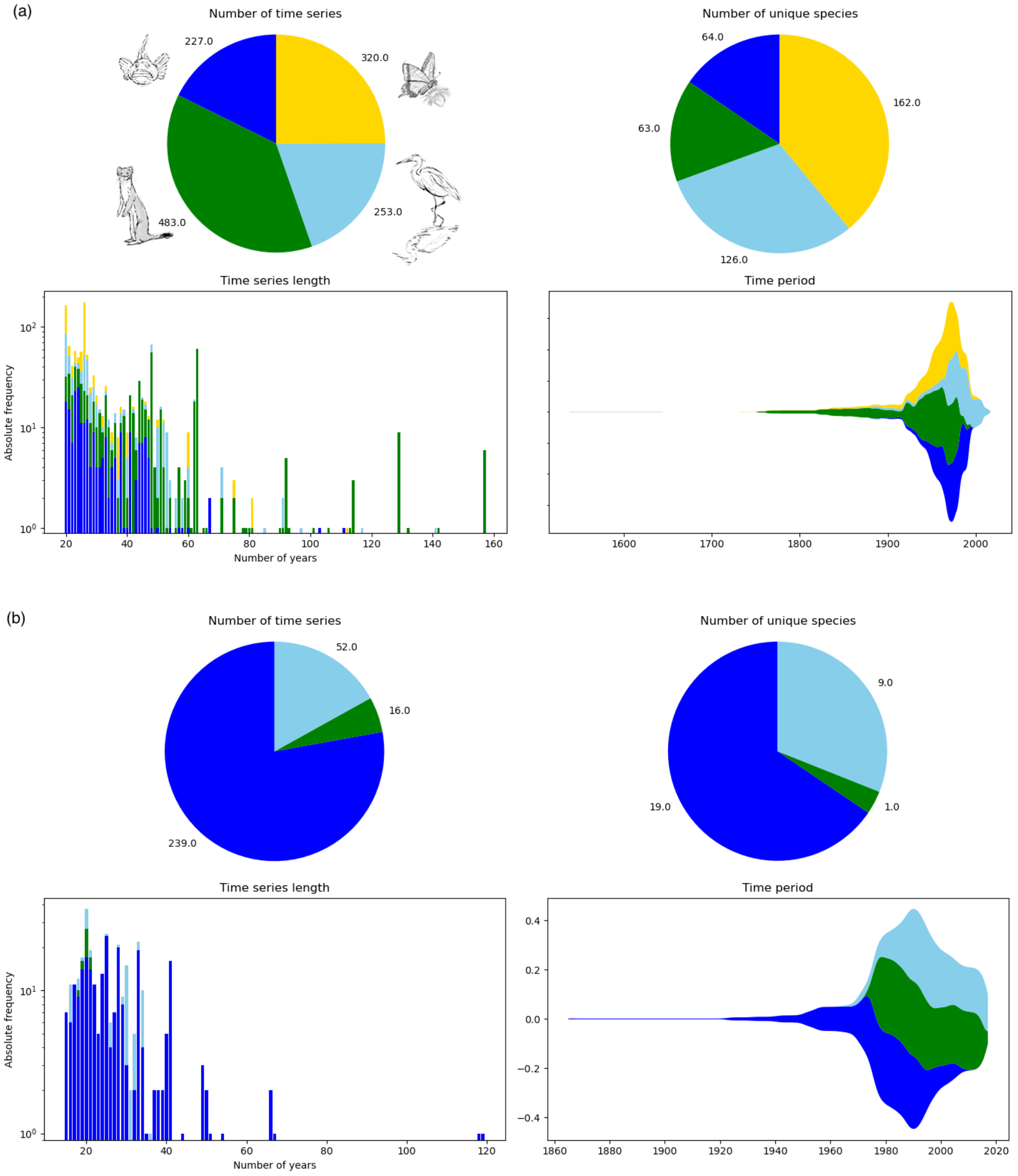


**Fig. S3. Summary of the analyzed ecological and phenotypic time series. (a)** The upper part of the figure summarizes the population time series characteristics analyzed for section §3 (number of time series, number of unique species, time series length, and time period), all characteristics sorted by taxonomic group. **(b)** The lower part of the figure summarizes the analogous results as above, for the phenotypic trait time series. Drawings: Andrea Klaiber, © Wildlife Analysis GmbH, Switzerland.

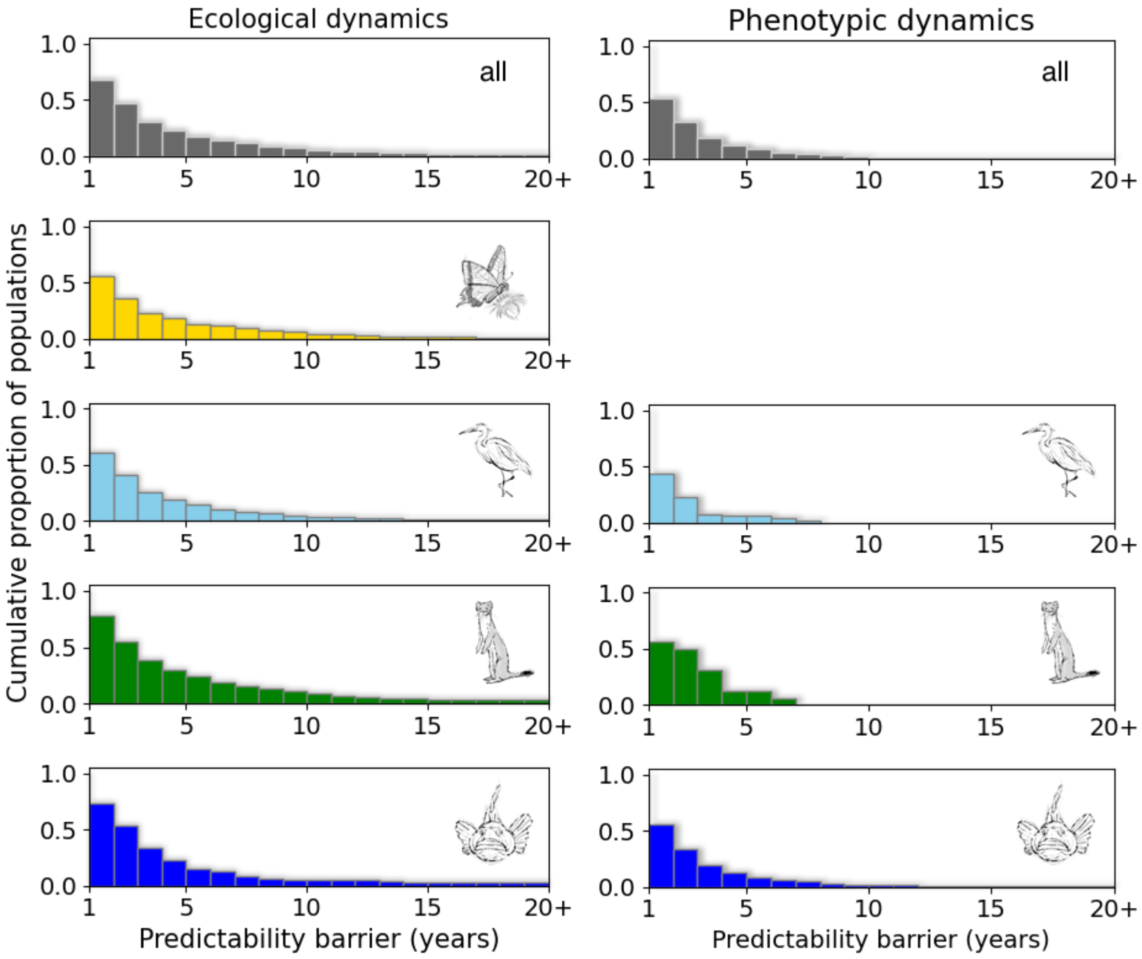


**Fig. S4. Predictability barrier.** The figure shows the predictability barriers (in years) of the 1283 analyzed invertebrate and vertebrate ecological time series (left column) and the 307 vertebrate phenotypic time series (right column). The first row gives the summary results for the respective data collection, while the following rows give the results by taxonomic group (insects, birds, mammals, and fish; cf. Fig. S3). In all panels the results are shown as the cumulative proportion of all respective populations being predictable at least $x$ years. Thus, one minus the proportion at a forecast horizon of one year (on the y-axis) indicates the proportion of population that is not even predictable one year ahead. Color-coding (taxonomic groups) as in Fig. S3. Drawings: Andrea Klaiber, © Wildlife Analysis GmbH, Switzerland.

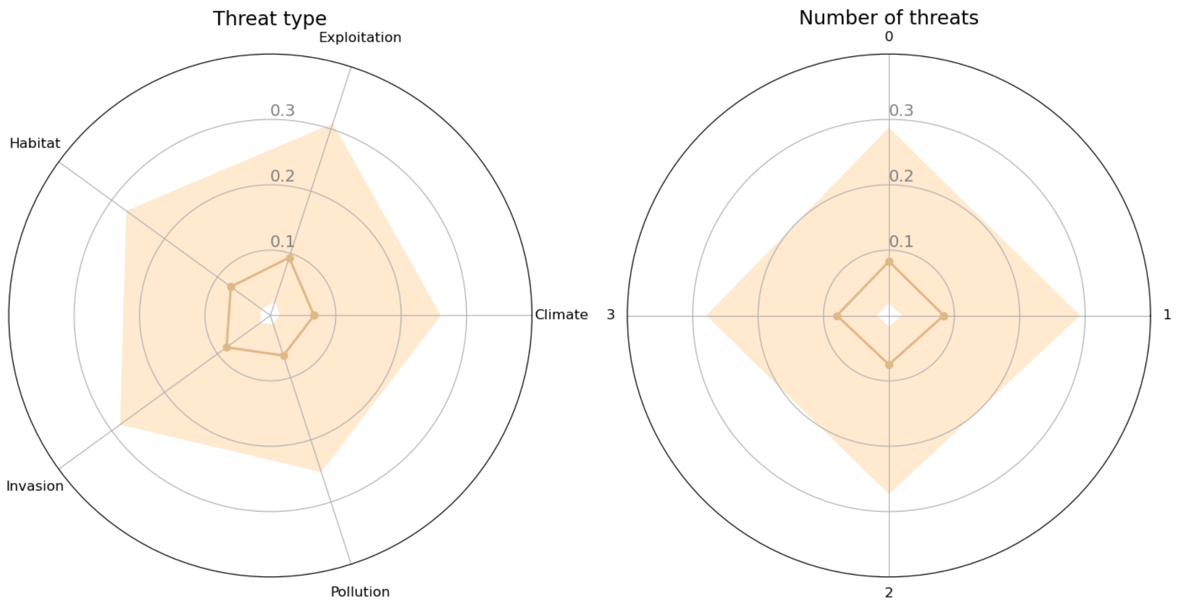


**Fig. S5. Predictability and human stressors (threats).** For the ecological time series, the two spider web plots show the intrinsic predictability results as measured by $PP(1)$ sorted by threat type (left) and total number of co-acting threats (right); see Appendix 4 for methodological details.

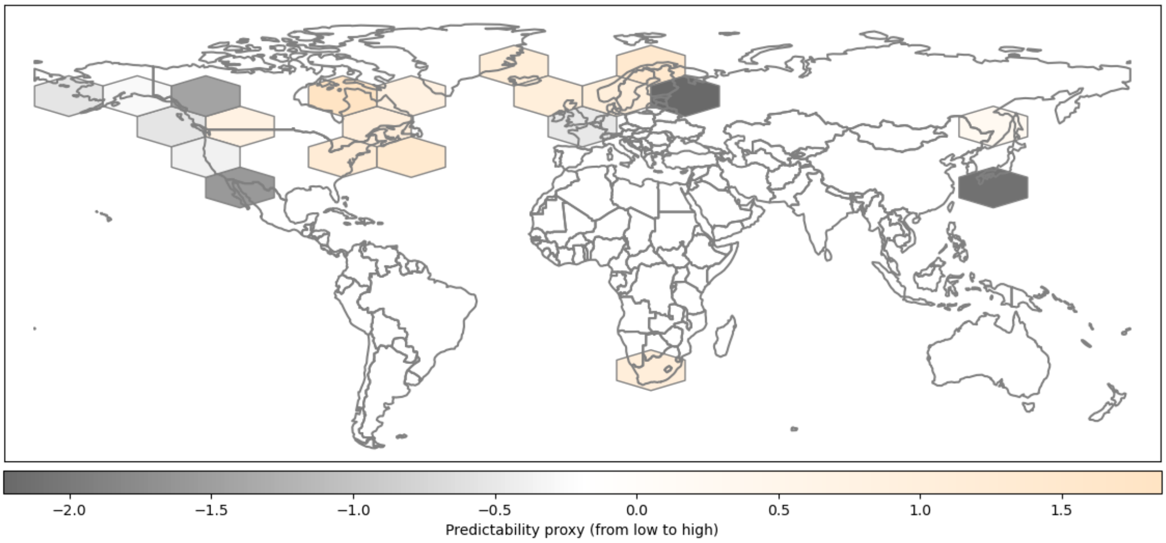


**Fig. S6. Geographic distribution of predictive power values (ecological dynamics): fish populations.** The world map depicts the analogous results to Fig. 4e, here however only using fish time series.

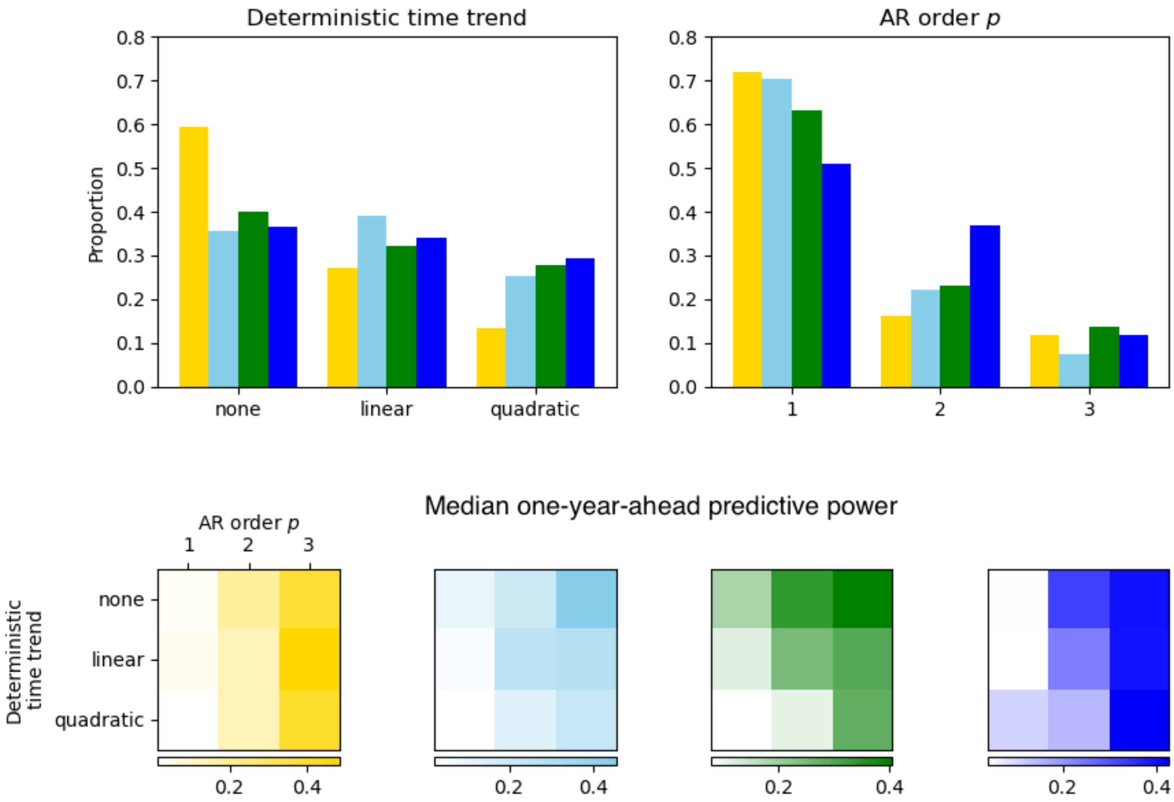


**Fig. S7. Time trend and model complexity measured by AR order *p* (ecological time series)**. The first row presents (left) the proportional distribution of statistically significant time trend types and (right) the distribution of the number of AR lags, both sorted by taxonomic group; color-coding as in Fig. S3. The second row gives, for the four taxonomic groups separately, the median one-year-ahead predictive power values, $PP(1)$ (color bar), depending on the time trend type and the number of AR lags.

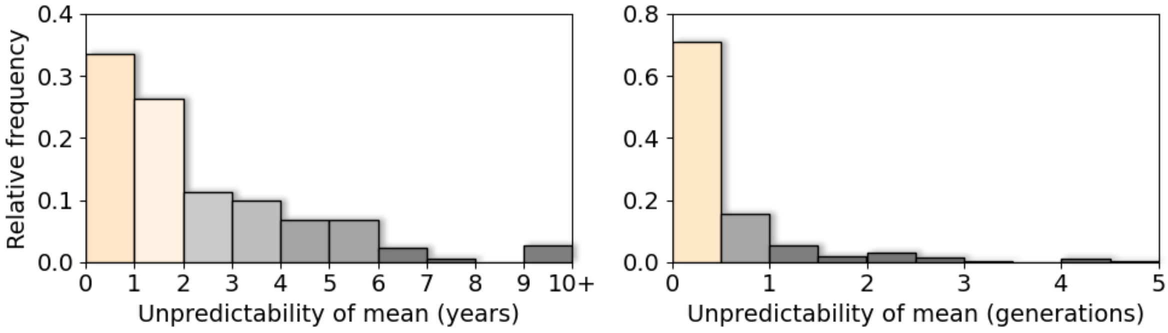


**Fig. S8. Forced predictability: phenotypic dynamics.** The left panel shows the same results as presented in Fig. 6e, while the right panel presents the analogous results expressed in terms of the number of unpredictable generations.

### Appendix 1: Computation of predictive power

#### 1.1 Without explanatory variables

The predictability measure predictive power, $PP(t)$, has been developed by Schneider and Griffies [4] to measure the uncertainty of a prediction, and the framework is derived using information-theoretic concepts, especially using the measure of entropy. The general form of $PP(t)$ can be re-formulated for $m$-dimensional Gaussian processes as

$$PP(t) \coloneqq 1 - \det\left(\frac{\mathbf{V}(t)}{\mathbf{V}_\infty}\right)^{1/(2m)}. \qquad \text{(eq. S1)}$$

Here, $\mathbf{V}(t)$ is the variance of the transition (also called forecast error) distribution and $\mathbf{V}_\infty$ is the variance of the stationary distribution of an $m$-dimensional Gaussian process, and $\det(\cdot)$ is the determinant. For univariate processes, eq. S1 reduces to eq. 1.1 in Box 1 (main text).

*Univariate time series*

The variance of the transition distribution and the variance of the stationary distribution of an univariate ARMA($p$,$q$) model – $\mathrm{v}(t)$ and $\mathrm{v}_\infty$ in eq. 1.1 (Box 1) – must first be calculated to compute the predictability measure predictive power, $PP(t)$. This can be done by recasting the ARMA($p$,$q$) model as a vector autoregressive process of order one, that is, a $(p+q)$-dimensional VAR(1) model. To this end, the ARMA($p$,$q$) model coefficients populate the two matrices $\mathbf{C}$ and $\mathbf{\Sigma}$, here exemplified using an ARMA(2,2) model:

$$\mathbf{C} = \begin{bmatrix} b_1 & b_2 & a_0 & a_1 \\ 1 & 0 & 0 & 0 \\ 0 & 0 & 0 & 0 \\ 0 & 0 & 1 & 0 \end{bmatrix}, \qquad \text{(eq. S2a)}$$

$$\mathbf{\Sigma} = \begin{bmatrix} \sigma^2 & 0 & \sigma^2 & 0 \\ 0 & 0 & 0 & 0 \\ \sigma^2 & 0 & \sigma^2 & 0 \\ 0 & 0 & 0 & 0 \end{bmatrix}, \qquad \text{(eq. S2b)}$$

where $\mathbf{\Sigma}$ is a covariance matrix, and $\sigma^2$ is the variance of the original univariate ARMA(2,2) process (Box 1, eq. 1.2b). The covariance matrix of the stationary distribution, $\mathbf{V}_\infty$, in vectorized form is defined as

$$\text{vec}\left(\mathbf{V}_{\infty(\text{VAR})}\right) := (\mathbf{I} - \mathbf{C} \otimes \mathbf{C})^{-\mathbf{1}}\text{vec}(\boldsymbol{\Sigma}), \tag{eq. S3}$$

where vec(·) is the vec-operator, $\mathbf{I}$ is an identity matrix of appropriate size, ⊗ denotes the Kronecker product, and subscript VAR refers to the recast model. To obtain the variance of the stationary distribution of the original univariate process ($\mathbf{V}_{\infty(\text{ARMA})}$, which is a scalar: $\text{v}_{\infty}$), the vector $\text{vec}\left(\mathbf{V}_{\infty(\text{VAR})}\right)$ is first reshaped as a matrix of size $(p+q) \times (p+q)$. Then, $\text{v}_{\infty}$ is given by the top-left scalar entry of this matrix. The covariance matrix of the transition distribution $t$ time steps ahead, $\mathbf{V}(t)$, in vectorized form is defined as

$$\text{vec}\left(\mathbf{V}(t)_{(\text{VAR})}\right) := (\mathbf{I} - (\mathbf{C} \otimes \mathbf{C})^{t})(\mathbf{I} - \mathbf{C} \otimes \mathbf{C})^{-1}\text{vec}(\boldsymbol{\Sigma}) \tag{eq. S4}$$

To obtain the variance of the stationary distribution of the original univariate process ($\mathbf{V}(t)_{(\text{ARMA})}$, which is a scalar: $\text{v}(t)$), again the vector $\text{vec}\left(\mathbf{V}(t)_{(\text{VAR})}\right)$ is first reshaped as a matrix of size $(p+q) \times (p+q)$, and the variance of the transition distribution for the forecast horizon $t$ time steps ahead is given by the top-left scalar entry of this matrix; for more examples and technical details, see Lütkepohl [5].

Predictive power in general has to be computed numerically, using the expressions given above (but see end of paragraph). The one-step-ahead predictive power, *PP*(1), nonetheless, for low-dimensional models can be computed directly using simple, analytically derived expressions. For example, for an ARMA(1,0) or ARMA(2,0) model

$$PP(1) = \begin{cases} 1 - \sqrt{1 - {b_1}^2} & \text{for ARMA(1,0)} \\ 1 - \sqrt{\dfrac{(1+b_2)\left(1 - {b_1}^2 + {b_2}^2 - 2b_2\right)}{1 - b_2}} & \text{for ARMA(2,0)} \end{cases}, \tag{eq. S5}$$

where $b_1$ and $b_2$ are the lag-1 and lag-2 autoregressive coefficients, respectively. As mentioned in Box 1 in the main text, notice that for univariate processes the process variance, $\sigma^2$, cancels out. For an AR(1) model, even a simple expression for $PP(t)$ can be derived, for any forecast horizon $t$: $PP(t) = 1 - \left(1 - {b_1}^{2t}\right)^{1/2}$. This expression allows to directly solving for the predictability barrier.

*Multivariate time series*

The computation of $PP(t)$ and the underlying variances for a multivariate dataset is analogous to the univariate case; in fact, the latter is a special case of the former. To construct the two matrices $\mathbf{C}$ and $\boldsymbol{\Sigma}$ (eq. S2a-b), every entry is now substituted by a matrix. For example, for a system of two variables (time series) instead of a single lag-1 coefficient $b_1$, we now use the matrix

$$\begin{bmatrix} b_1(1,1) & b_1(1,2) \\ b_1(2,1) & b_1(2,2) \end{bmatrix},$$

where for example $b_1(1,1)$ is the lag-1 coefficient of the first variable (as before) and $b_1(1,2)$ is the effect of the second variable (lagged by 1 time unit) on the first variable. For a multivariate case, the entry '1' in eq. S2a becomes an identity matrix, and the entry '0' in eq. S2a-b becomes a zero matrix, both of size $m \times m$. For the covariance matrix $\boldsymbol{\Sigma}$, every entry $\sigma^2$ now becomes a covariance matrix of size $m \times m$, and '0' again a zero matrix of size $m \times m$. The sizes of the resulting matrices $\mathbf{C}$ and $\boldsymbol{\Sigma}$ are then $m(p+q) \times m(p+q)$.

The computation of the stationary and transition variances follows the same procedure as detailed for the univariate case above. Note, however, that the vectorized variances (eq. S3-S4) will now have different sizes. Furthermore, after reshaping the vectorized variances as a $m(p+q) \times m(p+q)$ matrix, the covariance matrices $\mathbf{V}_\infty$ and $\mathbf{V}(t)$ are the top-left sub-matrix of size $m \times m$ of the respective matrix; here too, see Lütkepohl [5] for more examples and technical details.

##### 1.2 With explanatory variables

When future values of an explanatory variable $x(t)$ are assumed known, predictability as measured by $PP(t)$ depends on the transition and stationary distributions *around* the future mean values of $y(t)$ predicted by future $x(t)$. When the random errors are governed by an ARMA process (Box 1, eq. 1.2b), the formula for calculating $PP(t)$ using the residuals from a model including $x(t)$ is identical to the case when $x(t)$ is absent. Thus, the forecast can be decomposed into changes in the mean of the stationary distribution determined by future $x(t)$ and changes in the transition distribution relative to the stationary distribution given by $PP(t)$. To capture the information from $x(t)$ used for forecasts, we can define an overall predictive power as

$$PP(t)_{x(t)} := 1 - \frac{\sigma_\varepsilon}{\sigma_{y|x}}\big(1 - PP(t)\big), \qquad \text{(eq. S6)}$$

where $\sigma_\varepsilon$ is the standard deviation of the random errors of the fitted model to the observed $y(t)$ (Box 1, eq. 1.2b), and $\sigma_{y|x}$ is the standard deviation of the overall stationary distribution of $y(t)$ that depends on variation in $x(t)$. In this formulation of predictive power, $PP(t)_{x(t)} = 1 - \sigma_\varepsilon {\sigma_{y|x}}^{-1}$ when $PP(t) = 0$, which will be greater than zero provided $x(t)$ explains some of the variation in the observed values of $y(t)$; in this case the residual variance of the fitted model is less than the variance in $y(t)$ ($\sigma_\varepsilon^2 < \sigma_{y|x}^2$). A limitation of $PP(t)_{x(t)}$, however, is the assumption that $y(t)$ is stationary (so that $\sigma_{y|x}^2$ is finite) which requires that $x(t)$ be stationary. The covariate $x(t)$ might be non-stationary if, for example, $x(t)$ is climate temperature that is projected to constantly increase for the foreseeable future. In this case, $PP(t)_{x(t)}$ could still be used, but a value of $\sigma_{y|x}^2$ would have to be selected that makes sense for the specific question: for example, $\sigma_{y|x}^2$ could give the variance of the stationary distribution if $x(t)$ were to reach an equilibrium at some specific point in the future.

##### 1.3 Predictability of forced systems

As explained in the main text, intrinsic predictability as measured by predictive power, $PP(t)$, is only defined for stationary systems. Populations of concern are often non-stationary (e.g. trend-stationary), showing a time trend in abundance or phenotype [6]. DelSole and Tippett [7] have recently proposed a new measure of predictability suitable for forced systems, called total predictability, which encompasses the intrinsic predictability (they refer to it as initial-value predictability) and forced predictability (addressing a forced change in the mean of a process).

As illustrated in Fig. 6 in the main text, for a forced system three distributions are relevant: as for the case of unforced systems (Fig. 1), we have a stationary distribution (called climatological distribution in [7]) and a transition (forecast error) distribution; furthermore, we now also have the forced stationary distribution, which has a time-varying mean (see [7] for a changing variance). Note that the comparison of the forced stationary distribution to the current one does not mean that a time trend starts in the current year. Instead, this comparison is justified based on conditional independence: the authors' concept of generalized predictability can be understood as filtering out processes with a much longer time scale than the forecast horizon. To derive the total predictability measure, the authors start with the so-called mutual information, $M_{\mathrm{I}}$, which relates to predictive power as $PP(t) = 1 - \exp(-M_{\mathrm{I}})$; this is the intrinsic predictability of a stationary process (Box 1). Further, they show that total predictability of a forced system, $M_{\mathrm{T}}$, can be expressed as the sum $M_{\mathrm{T}} = M_{\mathrm{I}} + M_{\mathrm{F}}$, where $M_{\mathrm{F}}$ is the forced predictability due to a changing (forced) mean. Expressed in terms of predictive power, we have $PP(t)_{\mathrm{T}} = 1 - \exp(-M_{\mathrm{F}})\big(1 - PP(t)\big)$, which is eq. 2 in Box 3. When the variance of the unforced (i.e. 'original') and the forced stationary distributions are identical, the forced predictability takes the simple form $M_{\mathrm{F}} = \big(\mu(t) - \mu(0)\big)^2 (2\mathrm{v}_\infty)^{-1}$ (Box 3): here, $\mu(0)$ is the current mean from which predictions are made, $\mu(t)$ is the projected mean $t$ years ahead, and $\mathrm{v}_\infty$ is the variance of the stationary distribution of the univariate process (Box 1 and Appendix 1.1); the approach can be easily adapted to multivariate cases. Intuitively, bigger differences in the mean, $\mu(t) - \mu(0)$, will increase our ability to predict them. However, this is counteracted by the variance $\mathrm{v}_\infty$: more variable processes will make it harder to predict a changing mean. For further technical details and some worked examples based on forced AR(1) processes, we refer the reader to the original publication [7].

##### 1.4 Simulation study: AR(*p*) vs. ARMA(*p*,*q*)

To estimate the residual predictive power in the random errors of ARMA(*p*,*q*) models, decisions have to be made about what values to select for $p$ and $q$ in the fitted model. A sensible approach might be to fit models across a range of values of $p$ and $q$, and then select the best model according to a model selection criterion such as Akaike's Information Criterion (AIC). However, in our simulation study we found that setting $q$ = 0 and only considering AR($p$) models gave the most accurate estimates of the transition distribution which is the basic for the calculation of residual predictive power (Fig. S1).

We simulated datasets from ARMA(*p*,*q*) models (Box 1, eq. 1.2) for six sets of AR coefficients crossed with nine sets of MA coefficients to give 54 parameter combinations that generate distinct dynamics. For each of the 54 parameter combinations, we included a linear time trend by letting $\mu(t) = m_0 + m_1 t$ (Box 3), where the coefficient $m_1$ took values 0, –2.5, –5.0, –7.5, and –10.0. Finally, we considered time series of length 30 and 100. This led to a total of 540 scenarios for simulating time series. For each scenario, we simulated 1000 datasets of length 40 and 110. For each dataset, we estimated the ARMA(*p*,*q*) coefficients from the first 30 and 100 time points for $p$ = 1, 2 and 3, and $q$ = 0, 1 and 2 (nine combinations). We then computed the predicted transition distribution for the remaining 10 points in the time series. To assess the accuracy of the estimate of the transition distribution, for each simulated time series we scored whether the actual value fell within one standard deviation of the mean of the estimated transition distribution for times 1, 2, ..., 10 into the future past the time used to estimate the transition distribution. If the estimate of the transition distribution is accurate, then 68% of the projected values of the time series should fall within one standard deviation of the mean of the transition distribution.

The large number of parameter combinations (540) and fitting conditions (9) make it difficult to summarize all of the results. Nonetheless, the choice of setting $q = 0$ for the ARMA($p$,$q$) fitting was clear across all simulated conditions (Fig. S1). For the one-year-ahead transition distribution, fitting with $q = 1$ or 2 produced estimates of the transition distribution that were too narrow, so that a higher than expected proportion (0.32) of simulated points fell outside one standard deviation of the mean.

#### Appendix 2: Analysis of wolf population on Isle Royale

Given the illustrative nature of the analysis (main text section §2), the following models should be regarded as a basic starting point and not as the 'last word': for example, given the moderate population size of wolves, demographic stochasticity should in addition potentially be considered [8].

Wolves colonized Isle Royale, Lake Superior, from the nearby Canadian mainland around 1950. Research on this system now spans six decades, with monitoring starting in 1959. We limited the analyses to the years 1959 – 2011, after which the wolf population showed a heavy decline [9]. Moose make up the majority of the wolves' diet on Isle Royale, and this generates a tight dynamical coupling in this predator-prey system. Two additional factors have affected the wolf dynamics, a canine parvovirus (CPV) outbreak and inbreeding depression [10–12].

To compute predictive power for the wolf population (cf. Fig. 1), we used two models for the wolf dynamics, namely excluding or including covariates (Box 1). For both cases we log-transformed wolf and moose data (see below). For both cases we chose an autoregressive order $p = 1$, AR(1), based on previously published results [8]. Further, we chose a potential deterministic time trend function using AICc; see Appendix 3 for further details. According to model selection, the final models did not include a deterministic time trend.

For the analysis with covariates, we included moose data and an indicator variable summarizing the effect of CPV and inbreeding depression. For moose, although wolves show a clear preference for moose calves and senescent moose [8], given available data on senescent moose we included this prey stage group as covariate. Here, we used lagged data, so that the effect of moose in year $t - 1$ was expected to have an effect on wolves in year $t$. As for the second covariate, the CPV outbreak in 1980 took a considerable toll on the wolf population (cf. Fig. 1; [12]). Furthermore, it likely interacted with inbreeding depression so that the growth rate and consequently the average population size decreased [10–12]. We therefore included as a second covariate an indicator variable consisting of zeros up to the year 1980 and ones afterwards: the aim of this indicator variable therefore was to potentially adjust the population mean (Box 1, eq. 1.2a) after the CPV outbreak.

For predictions (cf. Fig. 1b), future values of the covariates were needed. For the indicator variable we continued setting it to one. For the senescent moose data, we fit an AR(1) model to them and used the predicted population mean as future values for the years after 2011.

#### Appendix 3: Analysis of Galápagos finch phenotypic time series

The dataset (see ESM Appendix 6) made available by Peter and Rosemary Grant [3] contains, among others, several long-term (1973-2012) phenotypic trait measurements of three Galápagos finch populations on Daphne Major Island: the medium ground finch (*Geospiza fortis*), the common cactus finch (*G. scandens*), and the large ground finch (*G. magnirostris*). The dataset also contains standard error data, but for the illustrative nature of our analyses in this paper we focused on yearly population means. Note that these error data could for example be used as measurement error variances when fitting an ARMA model in state-space form, using for example a Kalman filter. For our analyses, we used the three measurements beak length, depth, and width (Fig. S2). In addition to these time series, for all three analyzed species the dataset also contains the first component (PC1) from a principal component analysis applied to the three beak attributes, which we also analyzed (cf. Fig. 2 and Fig. S2).

For the univariate time series analyses we handled the data as described in Appendix 3, with two exceptions: first, we did not log-transform the PC1 time series; second, during model selection we only allowed for no time trend (i.e. a constant) or a linear time trend. The latter was motivated by the multivariate analyses (see below) which are generally more challenging in terms of fitting, and by restricting the time trend degree we thus restricted the maximum number of parameters; having the univariate and multivariate analyses have the same maximum degree makes the results comparable.

For the multivariate analyses we again handled the data as described in Appendix 3, without log-transforming the PC1 time series; we z-transformed the other time series separately. One challenge of fitting multivariate ARMA models, VARMA($p$,$q$), is the number of coefficients to be estimated in relation to the length and number of time series [5]. As an example, when analyzing three time series a VARMA(1,0) model will contain (i) three constants, (ii) potentially three linear time trend coefficients (iii) nine lag-1 coefficients, and (iv) six co-/variances (assuming a symmetric covariance matrix). Restricting the degree of time trend therefore results in three fewer parameters.

#### Appendix 4: Analysis of ecological and phenotypic time series collections

*Inclusion criteria and data handling*

To gain a bird's eye view on predictability across animal taxa, we analyzed a) ecological time series from the Global Population Dynamics Database (GPDD, [13]) and the InsectChange database (IC, [14]), and b) phenotypic time series from the PROCEED database [2]; see Appendix 6. The following criteria had all to be met in order for a time series to be considered for further analysis:

- only yearly data;
- no presence-absence data;
- a minimum length of 20 years (for ecological time series; because the available phenotypic time series were fewer, here we used a minimum length of 15 years), where the first and last entry was not a missing value;
- a maximum proportion of missing values of 25%;
- a maximum of 5 consecutive missing values.

We then log-transformed all time series that fulfilled the above criteria, except for proportional data that we logit-transformed. If the time series contained zeros, we first substituted them with missing values. Further, there were time series containing negative values: we did not log-transform these data, as they likely are indexed data. Finally, we z-transformed all log-transformed time series by subtracting the respective mean and dividing by the respective standard deviation.

*Model fit: model complexity and deterministic time trend*

For each time series ($n$ = 1283, Fig. S3), we used nine different models and selected the most appropriate one using Akaike's information criterion corrected for small sample size (AICc). The nine models differed in two aspects (Fig. S7):

- Model complexity: based on our simulation study (Appendix 1.4) we used three different autoregressive orders, AR($p$), $p$ = 1, 2, 3 (Box 1, eq. 1.2b), and we did not include moving average terms ($q$ = 0).
- Deterministic time trend: we tested three different versions, namely (i) no time trend, (ii) a linear one, or (iii) a quadratic one (i.e., a second-degree polynomial; Box 3).

*Explaining variation in PP(1): threats and geographic distribution*

Because we did not have population-specific threat data for the ecological time series collection, we proceeded as follows to investigate potential differences among threats or total number of threats in one-year-ahead predictive power values of ecological and phenotypic time series (Fig. S5). Using threat data from [1] for the taxonomic groups birds, fish, and mammals, we first computed the proportion of these groups within each threat, as well as for the total number of threats (Table S2). Subsequently, we logit-transformed all our $PP(1)$ values and for each threat or total number of threats computed a weighted mean of these values using the proportions in Table S2 as weights. Finally, we retransformed the resulting weighted means to the original scale.

To investigate potentially clustered geographic differences in predictability for the ecological time series (Fig. 4d), we proceeded as follows. After logit-transforming all $PP(1)$ values we first subtracted group-specific means of the four taxonomic groups. With the residuals we subsequently built the hexagonal grid shown in Fig. 4d. Using all analyzed populations (Fig. 4d) already shows that populations along the Pacific coasts in the northern hemisphere appear to be less predictable then populations in the North Atlantic. To better understand this pattern, we constructed additional hexagonal grid maps for every taxonomic group, which suggested that the pattern in Fig. 4d is driven by fish populations (Fig. S6). To understand this clustering, we used model complexity as measured by the AR order $p$ (Box 1) and found a highly significant association with the $PP(1)$ residuals used for Fig. 4d (Spearman's $r$ = 0.493, p-value < 0.0001), with a higher order expectedly generating a higher predictability (Fig. S7). Because the AR order $p$ captures the signature of strong biotic, trophic links of a population [15], the geographic predictability pattern in Fig. 4d indicates that the North Pacific populations harbor in their dynamics less effects of the food webs in which they are embedded as opposed to the North Atlantic.

### Appendix 5: Exploited populations

#### 5.1 Dynamical model of an exploited population

To investigate the potential effect of harvesting regimes on predictability (Box 2), we started with constructing a mechanistic discrete-time model for yearly population abundance data, $N_t$, suitable for harvested ungulate populations in temperate regions. The model includes a birth rate $\beta$, a natural mortality rate $\delta$, and density dependence with strength $\gamma$ where we assume that young-of-the-year will also affect it and that density dependence acts after autumn harvesting. The model further includes hunting bag, $H_t$, as a covariate and the random variable $\varepsilon_t$ capturing environmental variation,

$$N_t = N_{t-1}e^{\beta}\left(1 - \frac{H_{t-1}}{N_{t-1}e^{\beta}}\right)e^{-d-\gamma\ln\left(N_{t-1}e^{\beta}-H_{t-1}\right)+\varepsilon_t}. \qquad \text{(eq. S7)}$$

After log-transformation and simplification, the model reads

$$y_t = \varrho + (1-\gamma)y_{t-1} + (1-\gamma)\ln\left(1 - \frac{H_{t-1}}{N_{t-1}e^{\beta}}\right) + \varepsilon_t, \qquad \text{(eq. S8)}$$

where $y_t = \ln(N_t)$, and $\varrho = \beta(1-\gamma) - \delta$. We now express hunting bag $H(t)$ as a proportion of $N_t$, so that $H_t = pN_t$. Further, we define the proportion $p$ as a linear function of $y_t$, $p(y_t) = h_0 + h_1 y_t + \hbar_t$, where $\hbar_t$ are the residuals of the linear regression using data, i.e. $H_t {N_t}^{-1}$ vs. $y_t$. Thus, with $h_1 \equiv 0$ we have a (stochastic) proportion which does not depend on $y_t$. With $h_1 > 0$, the proportion harvested increases with increasing abundance. The case $h_1 < 0$, on the other hand, can be seen as a linear approximation to a constant yield system, that is, linearizing $p_t = Q_t {N_t}^{-1}$ at equilibrium, where $Q_t$ is the constant yield. Substituting $H_t = p(y_t)N_t$ in eq. Sx yields

$$y_t = \varrho + (1-\gamma)y_{t-1} + (1-\gamma)\ln\left(1 - (h_0 + h_1 y_{t-1} + \hbar_{t-1})e^{-\beta}\right) + \varepsilon_t. \qquad \text{(eq. S9)}$$

To make further progress we assume that the term $(h_0 + h_1 y_{t-1} + \hbar_{t-1})e^{-\beta}$ is not too big, say smaller than 0.2, so that $\ln(1-z) \approx \ln(\exp(-z)) = -z$. This approximation then helps simplifying eq. S9, leading to the final model

$$y_t \approx \mu_0 + c_1 \hbar_{t-1} + b_1 y_{t-1} + \varepsilon_t, \quad \text{(eq. S10)}$$

where $\mu_0 = \varrho - (1-\gamma)h_0 e^{-\beta}$, $c_1 = -(1-\gamma)e^{-\beta}$, and $b_1 = \left(1 - h_1 e^{-\beta}\right)(1-\gamma)$. This is the model reported in Box 2 as a dynamic regression model.

##### 5.2 Analysis of Alpine chamois populations

To investigate the potential effect of different harvesting regimes on predictability (Box 2), we analyzed 14 chamois populations at the Swiss administrative level of cantons (cf. Fig. 5a): Bern, Fribourg, Glarus, Jura, Luzern, Neuchâtel, Nidwalden, Obwalden, Schwyz, Solothurn, St. Gallen, Uri, Vaud, Zürich. We considered population dynamics at the level of Swiss cantons because it represents the appropriate level in terms of hunting legislation. For the chamois populations, data on yearly hunting bag as well as yearly abundance estimates are available [16] (data source: Appendix 6). For all populations, we used data for the years 1980 – 2020. For the population of St. Gallen, we substituted two data points (which we suspected to be outliers) with missing values. For all populations, the time series were potentially shortened in order for the first and last data points (hunting bag and abundance estimate) not to be missing values.

For every population we started with a linear regression to estimate the parameters for the (potentially) abundance-dependent proportion harvested, $p(y_t) = h_0 + h_1 y_t + \hbar_t$, where $p(y_t)$ and $y_t$ are based on the available data (see Appendix 5.1 above). Next, we fit eq. S10 supplemented with a potential deterministic time trend; see Box 2, eq. 2.2, and Appendix 3 for further details. The selected final model allowed computing predictive power for the harvested population. To compute predictive power for the unharvested population – think of it as a counterfactual – we used $b_1 = \left(1 - h_1 e^{-\beta}\right)(1-\gamma)$ to estimate the autoregressive parameter of the unharvested population, $(1-\gamma)$. Hereto, we set $\beta = 0.241$ based on previously published values [17]: after reproduction a population will have grown by 20-35%, and we used the mean of the log-transformed values.

#### Appendix 6: data sources

##### Section §2. What is predictability?

- Wolf and moose data [9]: https://isleroyalewolf.org/data/data/home.html (accessed 2022-10-25).
- Finch populations on Daphne Major Island [3]: https://datadryad.org/stash/dataset/doi:10.5061/dryad.g6g3h (accessed 2023-05-26).

##### Section §3. Predictability barriers across animal taxa

- Global Population Dynamics Database (GPDD) [13]: https://knb.ecoinformatics.org/view/doi:10.5063/F1BZ63Z8 (accessed 2022-04-01).
- Plot-level insect data (InsectChange database) [14]: http://onlinelibrary.wiley.com/doi/10.1002/ecy.3354/suppinfo (accessed 2022-11-28).
- Phenotypic trait time series [2]: https://github.com/photopidge/PROCEED (accessed 2023-07-03).

##### Section §4. Exploited populations

- Alpine chamois hunting bag and abundance data [16]: https://www.jagdstatistik.ch/de/home (accessed 2022-05-24).

##### Section §5. Predictability of forced systems

- See sections §2 and §3

#### References ESM


1. Jaureguiberry P *et al.* 2022 The direct drivers of recent global anthropogenic biodiversity loss. *Science Advances* **8**, eabm9982. (doi:10.1126/sciadv.abm9982)
2. PROCEED. 2022 Phenotypic Rates of Change Evolutionary and Ecological Database. See https://github.com/photopidge/PROCEED.
3. Grant PR, Grant BR. 2014 *40 Years of Evolution: Darwin's Finches on Daphne Major Island*. Princeton University Press.
4. Schneider T, Griffies SM. 1999 A Conceptual Framework for Predictability Studies. *Journal of Climate* **12**, 3133–3155. (doi:10.1175/1520-0442(1999)012<3133:ACFFPS>2.0.CO;2)
5. Lütkepohl H. 2005 *New Introduction to Multiple Time Series Analysis*. Springer. See https://link.springer.com/book/10.1007/978-3-540-27752-1.
6. Dornelas M *et al.* 2013 Quantifying temporal change in biodiversity: challenges and opportunities. *Proceedings of the Royal Society B: Biological Sciences* **280**, 20121931. (doi:10.1098/rspb.2012.1931)
7. DelSole T, Tippett MK. 2018 Predictability in a changing climate. *Clim Dyn* **51**, 531–545. (doi:10.1007/s00382-017-3939-8)
8. Vucetich JA, Peterson RO. 2004 The influence of prey consumption and demographic stochasticity on population growth rate of Isle Royale wolves Canis lupus. *Oikos* **107**, 309–320. (doi:10.1111/j.0030-1299.2004.13483.x)
9. Vucetich J, Peterson R. 2023 The Wolves and Moose of Isle Royale. See https://isleroyalewolf.org/ (accessed on 24 April 2023).
10. Adams JR, Vucetich LM, Hedrick PW, Peterson RO, Vucetich JA. 2011 Genomic sweep and potential genetic rescue during limiting environmental conditions in an isolated wolf population. *Proceedings of the Royal Society B: Biological Sciences* **278**, 3336–3344. (doi:10.1098/rspb.2011.0261)
11. Bozzuto C, Ives A. 2020 Inbreeding depression and the detection of changes in the intrinsic rate of increase from time series. (doi:10.13140/RG.2.2.23514.57289/1)
12. Wilmers CC, Post E, Peterson RO, Vucetich JA. 2006 Predator disease out-break modulates top-down, bottom-up and climatic effects on herbivore population dynamics. *Ecology Letters* **9**, 383–389. (doi:10.1111/j.1461-0248.2006.00890.x)
13. Prendergast J, Bazeley-White E, Smith O, Lawton J, Inchausti P, Kidd D, Knight S. 2010 The Global Population Dynamics Database. (doi:10.5063/F1BZ63Z8)
14. van Klink R *et al.* 2021 InsectChange: a global database of temporal changes in insect and arachnid assemblages. *Ecology* **102**, e03354. (doi:10.1002/ecy.3354)
15. Royama T. 1992 *Analytical Population Dynamics*. Springer Netherlands. (doi:10.1007/978-94-011-2916-9)
16. Federal Office for the Environment. 2023 Swiss federal hunting statistics. See https://www.jagdstatistik.ch/de/home (accessed on 19 April 2023).
17. Zeiler H. 2012 *Gams*. Österr. Jagd- u. Fischerei-Verlag.
